# The interaction between neural populations: Additive versus diffusive coupling

**DOI:** 10.1101/2021.11.29.470398

**Authors:** Marinho A. Lopes, Khalid Hamandi, Jiaxiang Zhang, Jennifer L. Creaser

## Abstract

Models of networks of populations of neurons commonly assume that the interactions between neural populations are via *additive* or *diffusive* coupling. When using the additive coupling, a population’s activity is affected by the sum of the activities of neighbouring populations. In contrast, when using the diffusive coupling a neural population is affected by the sum of the differences between its activity and the activity of its neighbours. These two coupling functions have been used interchangeably for similar applications. Here, we show that the choice of coupling can lead to strikingly different brain network dynamics. We focus on a model of seizure transitions that has been used both with additive and diffusive coupling in the literature. We consider networks with two and three nodes, and large random and scale-free networks with 64 nodes. We further assess functional networks inferred from magnetoencephalography (MEG) from people with epilepsy and healthy controls. To characterize the seizure dynamics on these networks, we use the escape time, the brain network ictogenicity (BNI) and the node ictogenicity (NI), which are measures of the network’s global and local ability to generate seizures. Our main result is that the level of ictogenicity of a network is strongly dependent on the coupling function. We find that people with epilepsy have higher additive BNI than controls, as hypothesized, while the diffusive BNI provides the opposite result. Moreover, individual nodes that are more likely to drive seizures with one type of coupling are more likely to prevent seizures with the other coupling function. Our results on the MEG networks and evidence from the literature suggest that the additive coupling may be a better modelling choice than the diffusive coupling, at least for BNI and NI studies. Thus, we highlight the need to motivate and validate the choice of coupling in future studies.

**Author summary:** Most models of brain dynamics assume that distinct brain regions interact in either an additive or a diffusive way. With additive coupling, each brain region sums incoming signals. In contrast, with diffusive coupling, each region sums the differences between its own signal and incoming signals. Although they are different, these two couplings have been used for very similar applications, particularly within models of epilepsy. Here we assessed the effect of this choice on seizure behaviour. Using a model of seizures and both artificial and real brain networks, we showed that the coupling choice can lead to very different seizure dynamics. We found that networks that are more prone to seizures using one coupling, are less prone to them using the other. Likewise, individual brain regions that are more likely to drive seizures when using additive coupling, are more likely to prevent them when using diffusive coupling. Using real brain networks, we found that the additive coupling predicted higher seizure propensity in people with epilepsy compared to healthy controls, whereas the diffusive coupling did not. Our results highlight the need to justify the choice of coupling used and show that the additive coupling may be a better option in some applications.

## Introduction

Modelling large-scale brain activity is key to better understanding macroscopic brain dynamics [1]. Merging such models and experimental data enables posing and testing hypotheses about brain function and dysfunction [1, 2]. There are two main classes of large-scale brain dynamic models, neural field models and brain network models (BNM) [1]. Neural field models treat the cortex as a continuous medium, whereas BNMs discretise the cortex into nodes. A node may typically represent a local population of excitatory and inhibitory neurons, whose activity may be modelled using a neural mass model such as the Wilson-Cowan model [3]. An ensemble of such coupled nodes is a BNM. The BNMs are particularly suited to study the role of brain network connectivity in shaping healthy and pathological dynamics as they can readily incorporate a brain connectome into a brain dynamics modelling framework [4]. For example, Hansen *et al*. [5] used a BNM and structural brain connectivity to simulate functional connectivity dynamics. Goodfellow *et al*. [6] used a BNM and functional connectivity derived from intracranial electroencephalography (EEG) to simulate and predict the outcome of epilepsy surgery. Demirtaş *et al*. [7] used a BNM to investigate the mechanisms responsible for connectivity changes in Alzheimer’s disease. Given the potential of BNMs to bring mechanistic understanding into the field of network neuroscience, it is important to be aware of its assumptions and choices.

BNMs may differ with regards to three main choices. First is the connectivity structure or network topology. BNMs may be used to investigate different types of networks, namely structural networks [8], or functional networks [6], which are inferred from different data modalities. Second is the choice of model to be employed to simulate the node dynamics. There is a wide range of model choices from biophysically realistic to purely phenomenological. For example, the Wendling model [6, 9] (a biophysical model), the Epileptor model [10] and the the subcritical Hopf bifurcation model [11] (two phenomenological models) have all been used within the context of modelling brain dynamics in epilepsy. Third is the interaction between nodes, which may be coupled with each other in a variety of ways. We identify two common coupling functions: *additive coupling* and *diffusive coupling*. For additive coupling the input to a node is a function of the sum of the activities of its neighbours. In contrast, for diffusive coupling the input to a node is a function of the sum of the differences between its activity and the activities of its neighbours. From a biophysical perspective, the additive coupling may be chosen when node activities represent currents, whereas the diffusive coupling may be appropriate if node activities represent electrical potentials. Even though these coupling definitions are different, they have been used for similar purposes in the literature. For example, in the epilepsy literature, the additive coupling has been used in studies to distinguish functional networks from healthy people and people with epilepsy [12], to simulate epilepsy surgery [6, 13, 14], and to model seizure propagation [15]. On the other hand, the diffusive coupling has also been used in studies to differentiate controls from people with epilepsy [11], to investigate epilepsy surgery [10, 16, 17], and to understand patterns of seizure emergence [18].

Although the additive and diffusive coupling frameworks have been used for such similar purposes, there has been no systematic assessment of the potential impact of this modelling choice on the resulting network dynamics and subsequent predictions. In this paper, we investigate whether this choice has an impact on network dynamics and model predictions in the context of epilepsy research. To this end, we focus on a phenomenological bi-stable model of seizure transitions that has been used with both additive and diffusive coupling in the literature [11, 12]. We test whether the additive and diffusive couplings lead to similar observations of the transient dynamics and predictions of seizure likelihood. Specifically, we use three salient measures, namely, escape times [11, 19], brain node ictogenicity (BNI) [6, 12], and node ictogenicity (NI) [6, 14]. The escape time quantifies the average time taken to transit from a resting state to a seizure state; the BNI measures the likelihood of a network to generate seizures; and the NI quantifies the contribution of single nodes to the network’s seizure propensity. We first apply all three measures to quantify the behaviour of artificial networks consisting of two, three and 64 nodes with additive and diffusive coupling. We then test whether the two couplings provide similar results in terms of BNI when applied to functional brain networks inferred from resting-state magnetoencephalography (MEG) with the aim of distinguishing individuals with juvenile myoclonic epilepsy (JME) and healthy controls. Concordant results in terms of escape time, BNI and NI between models using additive and diffusive coupling would imply that the choice of coupling is inconsequential with little impact on the predictions relevant to epilepsy; whereas discordant results would ask for careful consideration when choosing the coupling.

## Materials and methods

### Phenomenological model of seizure transitions

To assess seizure-like dynamics with a BNM using both additive and diffusive coupling, we consider a commonly used phenomenological model of seizure transitions that is based on the normal form of the subcritical Hopf bifurcation [11–13, 16, 17, 19]. In this model, each network node can be represented by a noisy bi-stable oscillator, where a stable fixed point coexists with a stable limit cycle. Fluctuations around the fixed point represent resting dynamics, whereas large oscillations around the limit cycle correspond to seizure dynamics. Transitions between the two states are driven by noise and the influence of other nodes in the network.

The network dynamics is described by the following system of stochastic differential equations:

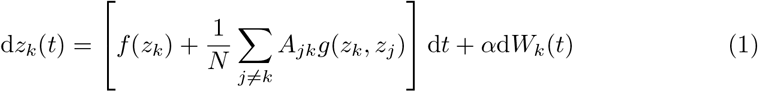

where *z_k_*(*t*) is a complex variable that describes the dynamics of node *k* (*k* = 1, 2, …, *N*, and *N* is the number of nodes). The function *f*(*z*) that defines the activity of a single node is,

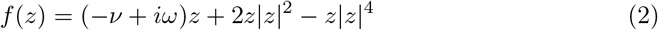

The parameter *v* controls the stability of the node and *ω* defines the frequency of the oscillations that the node may display depending on *v*. At *v* < 0, the origin is an unstable fixed point and the node oscillates around a stable limit cycle. At *v* = 0, the unstable point becomes stable in a subcritical Hopf bifurcation. For 0 < *v* < 1, the node is bi-stable with a stable fixed point at the origin and a stable limit cycle separated by an unstable limit cycle. At *v* = 1, the stable and unstable limit cycles meet each other in a saddle-node bifurcation. In this paper, we use *v* = 0.2 and we fix *ω* = 20 in line with previous studies [11, 17, 19, 20] (except in the two-node networks, where we use *ω* = 0, as described below). Each node has an independent (identically distributed) complex white noise process 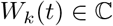, the strength of which is governed by the noise amplitude *α* > 0.

The interaction between nodes is determined by the adjacency matrix *A_jk_* and the coupling function *g*(*z_k_, z_j_*). Here we use three coupling functions:

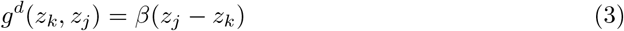

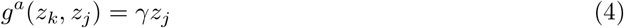

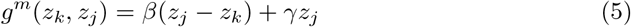

corresponding to diffusive coupling (3), additive coupling (4), and a combination of the two which we will call ‘mixed’ coupling (5). Note that if *β* = 0, then *g^m^*(*z_k_, z_j_*) = *g^a^*(*z_k_, z_j_*) = *γz_j_*, whereas if *γ* = 0, then *g^m^*(*z_k_, z_j_*) = *g^d^*(*z_k_, z_j_*) = *β*(*z_j_* – *z_k_*). The diffusive coupling was used for example by Benjamin *et al*. [11], Terry *et al*. [21], Hebbink *et al*. [16], Creaser *et al*. [18, 19], and Junges *et al*. [17]. The additive coupling was used for example by Petkov *et al*. [12], Sinha *et al*. [13], and Junges *et al*. [20]. To the best of our knowledge, the mixed coupling has never been considered.

### Quantifiers of seizure transitions

To quantify and compare the effect of the three coupling functions on the behaviour of the BNM we use the following measures. The first is based on escape time theory, and has previously been used to classify the behaviour of motif networks of this BNM with diffusive coupling [11, 18, 19]. The second is brain network ictogenicity which is suitable for comparing the propensity of different networks to generate seizure dynamics (ictogenicity) [6, 12, 14, 20]. The third is node ictogenicity, a quantity that assesses the contribution of each node to the network’s ictogenicity [6, 14, 20].

#### Escape time

We will first characterise the transition to seizure dynamics in two coupled nodes. To this end, we consider the mean time taken for nodes to transition from the stable fixed point to the stable oscillatory seizure state. For one node, we define the *escape time* λ as the moment *t* at which the amplitude of its activity *z* crosses a given threshold. This *escape threshold* is usually chosen to be the unstable point (or gate) between stable states. The initial condition of all nodes is the resting state, here the fixed point at the origin. We only consider transitions from the resting state to the oscillatory seizure state, as for our chosen parameter values the oscillatory stable state is much more strongly attracting than the fixed point, and so transitions back again to the resting state happen on a much longer timescale. The escape time λ is a random variable and so we define the mean escape time as 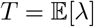. The mean escape time of a single node (1) for *k* =1 has been fully characterised using Eyring–Kramers’ escape time theory in [11, 19].

For two coupled nodes we define the first escape time as the time the first node transitions to the oscillatory state, which can be either node. The second escape time is then the time that it takes the other node to transition after the first one has escaped. As before these are random variables and so we can define the *mean first escape time* and *mean second escape time*; here we will refer to these means as the first escape and second escape. The first and second escape times in a two node system with diffusive coupling were characterised in [19]. Here, we will extend this work to compare the effect of all three coupling functions on the escape times.

We identify qualitatively different regimes of escape time behaviour. By converting the BNM with two nodes into polar coordinates *z_k_*(*t*) = *R_k_*(*t*) exp[*ιθ_k_*(*t*)] using Itô’s formula, we can restrict our attention to the amplitude dynamics of the system. Note that the oscillatory phase of the periodic orbit does not affect the escape times as we consider the case where the phase difference between nodes is zero. The steady states of the amplitude dynamics are the ”transition states” where either none, one or both nodes have escaped. The number and stability of these states change with the strength of the coupling. We perform bifurcation analysis on the transition states as we vary each of the coupling strength parameters *β* and *γ*, using specialist continuation software AUTO-07P [22]. We consider here only the symmetrically coupled case (bi-directional coupling) for which we can consider the amplitude dynamics as a potential system and identify the potential landscape *V* [19].

We numerically compute the mean escape times for two bidirectionally coupled nodes with each of the three coupling functions using custom code written in MATLAB. For each coupling strength we compute 1000 simulations of the two node model using the stochastic Euler-Maruyama method with step size *h* = 10^−3^ and initial conditions *z_k_*(0) = 0. To leading order the escape times do not depend on the choice of escape threshold provided it lies beyond the unstable limit cycles, and in the following we fix it to be a node amplitude of 0.5 in line with [11, 19]. For each simulation we identify the first and second escape time, then take the means over the 1000 simulations. The escape times do not depend on *ω* and so we set this to zero in our simulations.

#### Brain network ictogenicity

To characterise seizure-like dynamics in networks, we use the concept of brain network ictogenicity (BNI) [6, 12, 14, 20]. The BNI quantifies the likelihood of a network to generate seizures and corresponds to the average time that each network node spends in the seizure state. Since the initial conditions are such that all nodes start in the resting state and once they transition to the seizure state, they do not return to the resting state, then the BNI is formulated as

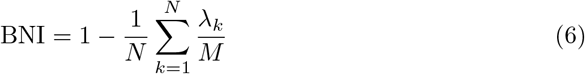

where *M* is a sufficiently long simulation time and λ_*k*_ is the escape time of the *k*^th^ node. The higher the fraction λ_*k*_/*M* is, the longer the node *k* takes to seize. If node *k* does not escape during the simulation time, we take λ_*k*_ = *M*. Thus, the BNI ranges from 0 to 1, where networks with low BNI have more nodes with high escape times, whereas networks with high BNI have more nodes with low escape times. This definition of BNI is equivalent to the seizure likelihood measure used by Sinha *et al*. [13]. Whilst we use the escape times to study two interacting nodes, we compute the BNI for three-node motifs and larger networks (see section). To compute the BNI, we integrated the stochastic equations (1) using the Euler-Maruyama method with step size *h* = 10^−3^, set initial conditions *z_k_*(0) = 0, fixed *M* = 50/*h*, and averaged the BNI across 1000 noise realisations.

#### Node ictogenicity

To quantify the contribution of each node to the network’s ability to generate seizures, we use the concept of node ictogenicity (NI) [6, 14, 23]. The NI^(*k*)^ measures the relative difference in BNI upon removing node *k* from a network:

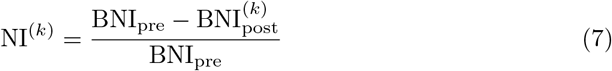

where BNI_pre_ is the BNI prior to node removal, and 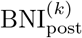 is the BNI after the removal of node *k*. Note that to compute 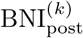, the coupling term in (1) is not normalised by *N*, but by *N* +1 (i.e., the size of the network before node removal), so that the effective coupling strength is kept fixed. As in previous studies, we set the coupling strength parameters such that BNI_pre_ = 0.5 [6, 14, 23] (except in the three node networks, where we use the network BNI computed as above), and the same parameters are used to compute 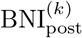. The NI^(*k*)^ ranges between −1 and 1, where NI^(*k*)^ = –1 means that removing node *k* increases the ictogenicity of the network to its maximum 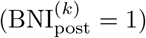; NI^(*k*)^ = 0 means that the removal of node *k* has no impact on the ictogenicity of the network 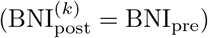; and NI^(*k*)^ = 1 means that removing node *k* stops all seizure activity in the network 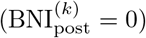. Here we use it to assess whether the relative importance of nodes for the network ictogenicity depends on the coupling function.

To compare different NI distributions obtained using the different coupling functions, we consider the weighted Kendall’s rank correlation *τ* [14, 23–25],

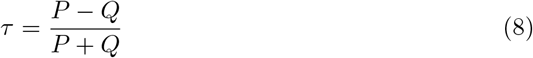

where *P* is the number of pairs of nodes with the same order in two rankings (e.g., pairs of NI values using the additive coupling that are ordered in the same way as pairs of NI values using the diffusive coupling), and *Q* is the number of pairs in reverse order. To account for the relative NI values in the comparison between distributions A and B, the contributions to *P* and *Q* are weighted by the product of the distances in NI in the two pairs, 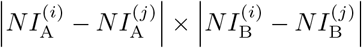, where 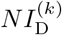 is the ictogenicity of node *k* in distribution D. The value of *τ* ranges from −1, i.e., all pairs in reverse order, to 1, i.e., all pairs with the same order.

### Artificial networks

To better characterise differences and similarities between the different coupling choices, we investigated the model in a variety of networks. This was to ensure that our observations were not specific to a certain kind of network structure. Indeed, we have previously shown that the *BNI* and *NI* are a function of the network topology [14]. Thus, besides analysing the dynamics of two and three interacting nodes, we also simulated large networks with *N* = 64 nodes. This is a typical network size in studies with BNMs applied to functional brain networks [6, 13, 14]. We considered random and scale-free networks, both directed and undirected [14, 26, 27]. We generated random networks using the Brain Connectivity Toolbox [28]. To build undirected scale-free networks with degree distribution *P*(*x*) ∝ *x^−a^*, we used the static model [29] and *a* = 3. Finally, we employed the Barabási-Albert algorithm to construct directed scale-free networks [30]. We considered networks with mean degree *c* =4 and *c* =8. In the case of directed networks, we used mean in-degree *c*_in_ equal to the mean out-degree *c*_out_, *c*_in_ = *c*_out_ = *c*. We discarded networks with disconnected components and analysed 10 network realisations per network topology. Thus, we studied 80 networks.

### MEG functional networks

To verify how our results generalise to real-world brain networks we compared additive and diffusive coupling in terms of BNI on MEG functional networks. We used resting-state MEG functional networks from people with JME and healthy controls. This MEG dataset was previously used to demonstrate that the BNI framework was capable of differentiating the two groups of individuals [31]. In that study, Lopes et al. [31] used the theta model with additive coupling (see the Supplementary Material for a description about the theta model), and showed that individuals with epilepsy had higher BNI than controls as hypothesized. The difference here is that we use the bi-stable model instead of the theta model, and that we consider the MEG networks specifically to compare the effect of additive versus diffusive coupling.

We refer the reader to Lopes et al. [31] for details about the participants, MEG acquisition, pre-processing, source mapping, and functional network construction. Briefly, the dataset comprises 26 people with JME and 26 controls. The control group was age and gender matched to the JME group (the median age was 27 and there were 7 males in both groups). This study was approved by the South East Wales NHS ethics committee, Cardiff and Vale Research and Development committees, and Cardiff University School of Psychology Research Ethics Committee. Written informed consent was obtained from all participants.

Approximately 5 minutes of resting-state MEG data were acquired using a 275-channel CTF radial gradiometer system (CTF System, Canada) at a sampling rate of 600 Hz. The first 200 s of artefact-free data were selected for each individual. The data were then filtered in the alpha band (8–13 Hz) and down-sampled to 250 Hz. Subsequently, the underlying sources were inferred using a linear constrained minimum variance (LCMV) beamformer on a 6-mm template with a local-spheres forward model in Fieldtrip [32]. The source signals were then mapped into the 90 brain regions of the Automated Anatomical Label (AAL) atlas [33].

To obtain MEG functional networks, the source reconstructed data were divided into 10, non-overlapping, 20 s segments. A functional network was computed from each segment using a surrogate-corrected amplitude envelope correlation (AEC) with orthogonalised signals [31, 33]. Thus, we considered 10 MEG functional networks per individual. We then took the average of the 10 networks and analysed one average network for each individual. To then measure BNI on these networks, we computed seizure-like dynamics using (1), with *A_jk_* equal to the average surrogate-corrected AEC values of the functional networks.

## Results

### Two coupled nodes

To illustrate fundamental differences between the coupling functions we first consider the simplest case of two bidirectionally coupled nodes. Figure 1 shows how each coupling function changes the transient behaviour of the two node system. The bifurcation diagrams of the amplitude of the transitions states (equilibria) *R*_1_ of node 1 are plotted for the additive and diffusive coupling functions, (4) and (3) respectively. Due to the symmetry of the system the diagrams plotted against *R*_2_ are identical. For each coupling type the transition states undergo saddle node and pitchfork bifurcations as the coupling strength increases. We follow these bifurcation points in both *β* and *γ*. The resulting two dimensional bifurcation diagram shows that these bifurcations delineate qualitatively different dynamic regimes. Example simulations of each regime are illustrated on the potential landscape with the locations of the transition states (equilibria).

**Fig 1.**
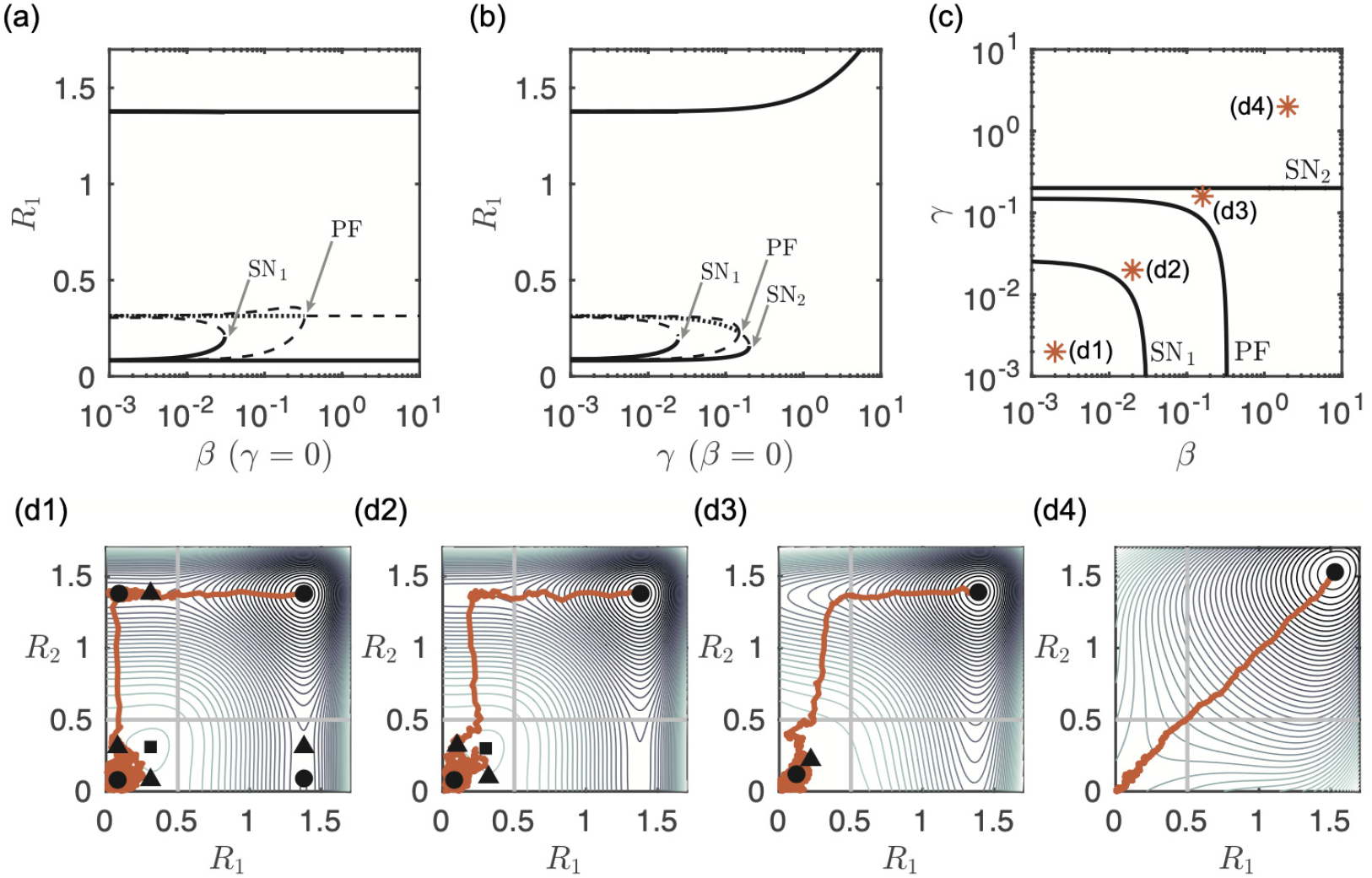
Bifurcation diagrams and example simulations of the bidirectionally-coupled two-node system. Shown are the bifurcation diagrams of amplitude *R*_1_ of the transition states (equilibria) as a function of *γ* for *β* = 0 (additive coupling) in panel (a), and of *β* for *γ* = 0 (diffusive coupling) in panel (b). Stable states are shown as solid lines, saddle states as dashed lines and unstable as dotted lines. Saddle node bifurcations are labelled SN_1_ and SN_2_, the pitchfork bifurcation is labelled PF. Panel (c) shows the position of the saddle node and pitchfork bifurcations in the (*β, γ*)-plane. The orange stars indicate the values for which example simulations are shown in panel (d). Each subpanel (d) shows an example simulation in orange starting at *z_k_*(0) = 0 with the steady states in black (the circles are stable, the triangles are saddle and the squares are unstable (source)). Contour lines of the potential landscape are also plotted in shades of grey. The escape threshold for each node is shown as a solid line at 0.5.

The bifurcation diagrams show that when the nodes are uncoupled, *β* = *γ* = 0, there are 9 transition states of the system. These corresponding to all possible pairs of the states of the individual nodes, resting state (not escaped), seizure state (escaped) and the threshold state. When the coupling strength is weak (*β* = *γ* < 0.01) the nodes behave as if uncoupled and all 9 states persist. As each coupling strength is increased the equilibria first undergo a saddle node bifurcation where the partially escaped state (node 1 has escaped but node 2 has not, and vice versa) disappear and are no longer attractors of the system. However, due to the contours of the potential landscape (panel (d2)), realisations spend some time in the vicinity of the partially escaped state. This means that while escape of both nodes is inevitable there is a delay between the first and second escapes. The unstable equilibrium then undergoes a pitchfork bifurcation where the system becomes synchronous in the sense that escapes from the resting to the seizure state for both nodes occur in quick succession. The key difference in dynamics comes when the additive coupling induces a further saddle node bifurcation and the only attractor of the system is the state in which both nodes have escaped to the seizure state. This regime never occurs with diffusive coupling only. Essentially, for a sufficiently large additive coupling strength, both nodes are forced immediately into the seizure state. In contrast, for large diffusive coupling strength the nodes stay in their starting resting state until noise eventually kicks both of them simultaneously into the seizure state.

Figure 2 shows the first and second escape times for different values of *β* and *γ*. The mean first escape times for the diffusive only coupling (*γ* = 0) and the additive only coupling (*β* = 0) show opposite trends. Note that we do not distinguish between node 1 or node 2 escaping first. For diffusive coupling the escape times increase as *β* increases. The coupling has an inhibitory effect and the nodes spend longer in the resting state where neither has escaped. For additive coupling the escape times decrease as *γ* increases as the coupling has an excitatory effect. Figures 1(c) and (d4) show that for large *γ* only the fully escaped equilibrium remains and so the first escape time depends only on the starting position in the (*R*_1_, *R*_2_)-plane and level of noise. With the mixed coupling function (5), when the coupling is weak, *γ, β* < SN_1_, the system behaves as if uncoupled and neither type of coupling dominates. When *γ* is small (*γ* < 10^−1^), the first escape times follow the same pattern as for the diffusive only coupling and for large *β* the diffusive coupling dominates. Conversely, when *β* is small (*β* < 10^−1^), the first escape times follow the same pattern as for the additive only coupling and for large *γ* the additive coupling dominates.

**Fig 2.**
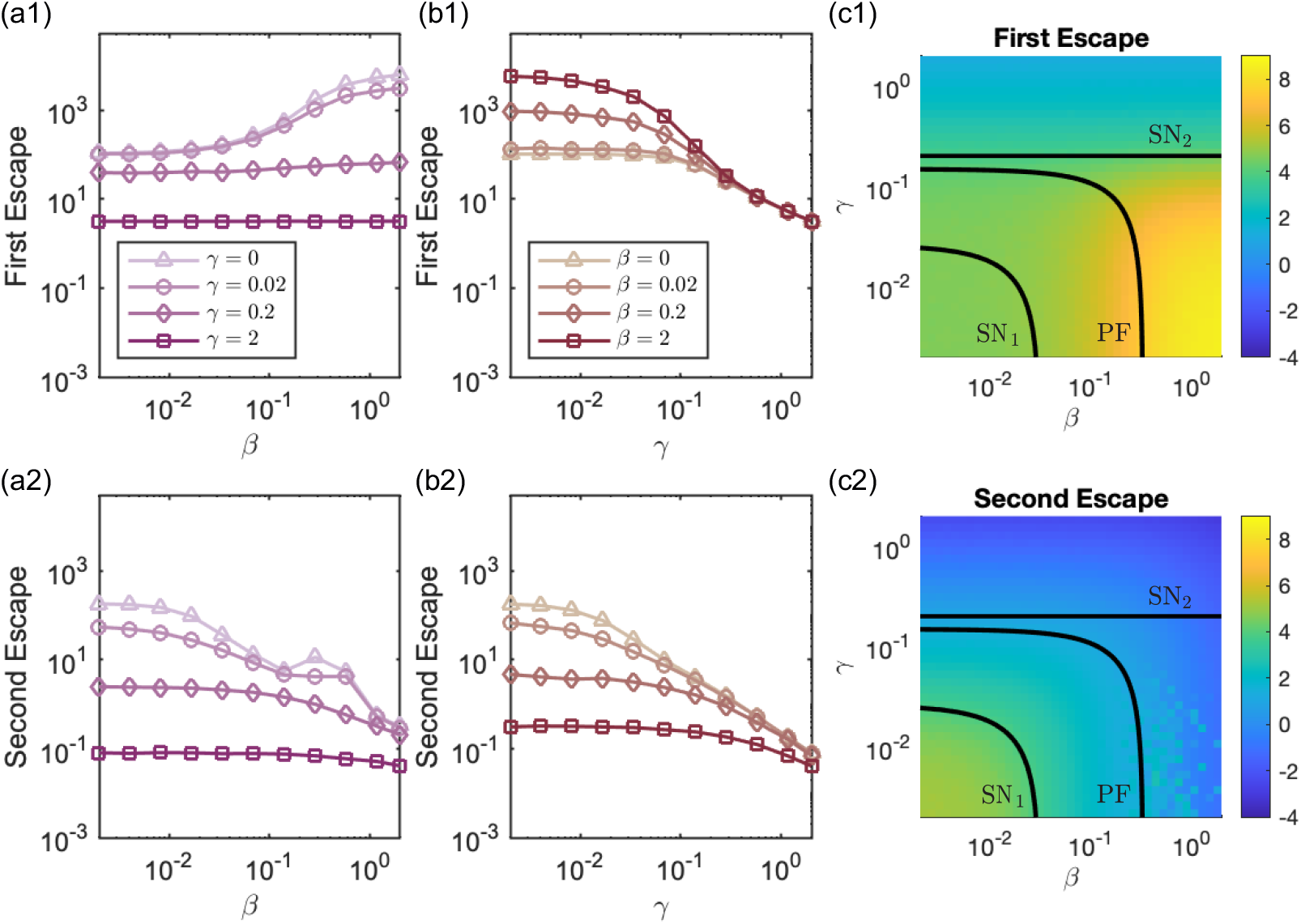
Mean times for the first node to escape (First Escape, row 1) and second node to escape (Second Escape, row 2). Column (a) shows mean escape times against *β* for different values of *γ*; Column (b) shows mean escape times against *γ* for various *β* values. A legend is given for each column. Column (c) plots the mean escape times on the (*γ,β*)-plane with the bifurcation curves overlaid.

The diffusive dominated coupling is characterised by the area of very large escape times in yellow for high *β* and low *γ*. However, for large *β* and *γ* > SN_2_ the first escape time no longer depends on *β* and additive coupling dominates. This is illustrated by the almost flat lines for *γ* = 0. 2 and 1 in panel (a1) and the coalescence of all the lines in panel (b1) for *γ* > 0.1. The mean second escape times show a decreasing trend with increasing coupling strength for both the additive and diffusive only cases. We note that around the pitchfork bifurcation (PF) in the diffusive case (large *β*) the second escape times become noise dominated.

Taken together, these escape times show that as coupling strength is increased the state where only one node has escaped disappears and the behaviour of the nodes synchronises. In other words, as soon as one node escapes the other immediately follows. This is because when either of the coupling strengths are large the input from connected nodes dominates the dynamics. The key difference is the time that it takes the first node to escape, which is fundamentally different depending on whether additive or diffusive coupling dominates the system.

### Three-node networks

To consider the effect of network structure (topology) on the escape times of the network and its nodes we consider the BNI and NI of three-node motifs. For simplicity, in this section we compare only additive and diffusive coupling. Figure 3 shows the BNI computed via (6) and NI computed via (7) for all non-isomorphic three-node networks. For each network, we observe that the BNI is higher for the additive coupling than for the diffusive coupling. Furthermore, we observe that networks with higher number of connections tend to have higher BNI for the additive coupling, but lower BNI for the diffusive coupling. These results show that, networks with more additive connections are self excitatory and have more nodes with low escape times, whereas networks with more diffusive connections are more self inhibitory and have more nodes with low escape times. It also indicates that increasing the number of connections has a similar effect as increasing the coupling strength. Thus, the discrepancy in BNI between the two couplings tends to be greater in networks with more connections.

**Fig 3.**
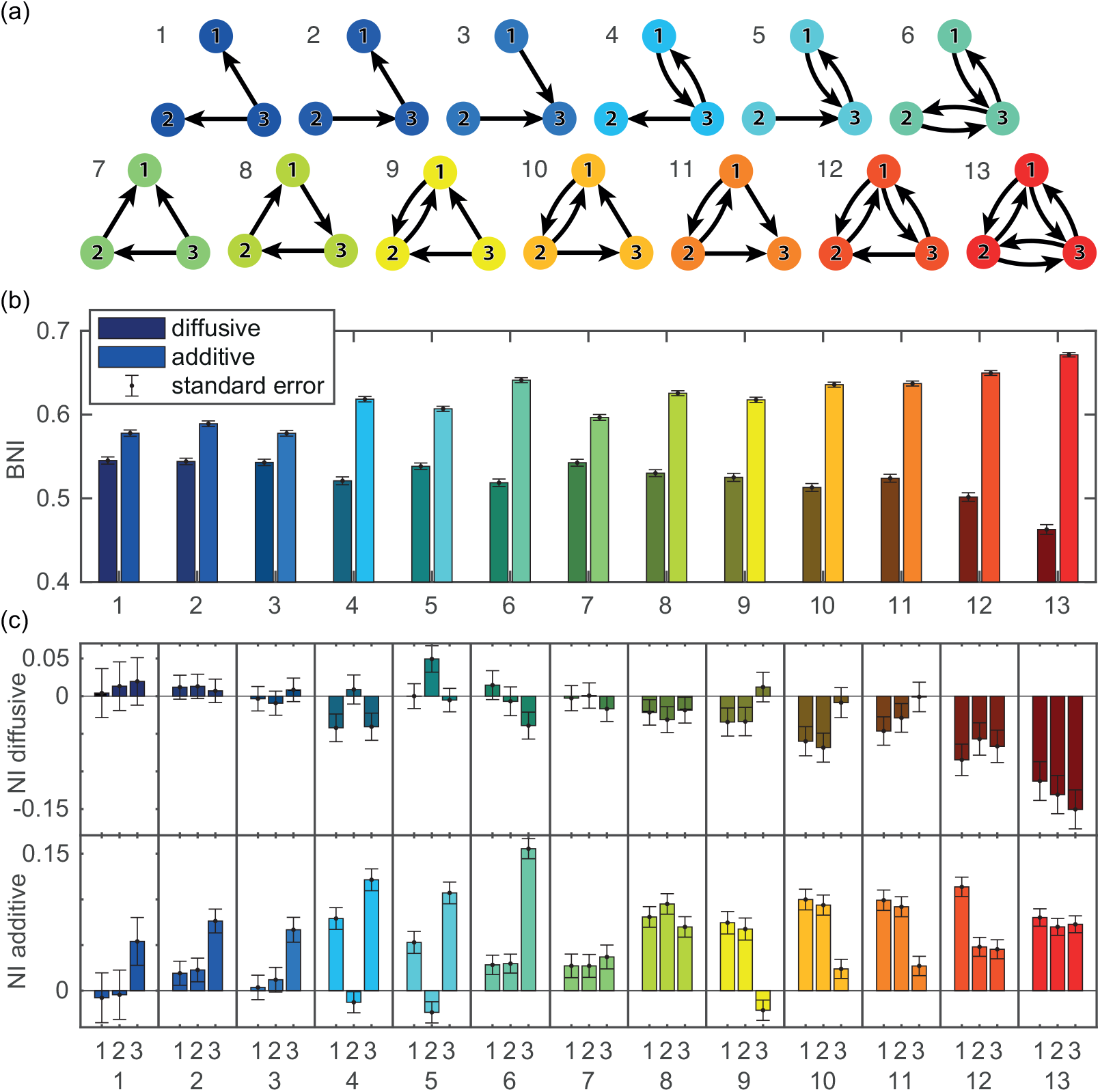
BNI and NI of all three-node networks for additive and diffusive couplings. Panel (a) shows the coupling structure for all 13 non-isomorphic three-node networks. Panel (b) shows the BNI for each network with either diffusive (darker bars) or additive coupling (lighter bars). Panel (c) shows the NI for each node (1–3) of each network (1–13) with either diffusive (top row, darker coloured) or additive (bottom row) coupling. The NI distributions were computed using as BNI_pre_ the BNI values in panel (b). We used *β* = 0.1 for the diffusive coupling, *γ* = 0.1 for the additive coupling, and *α* = 0.03 for both couplings. Standard error bars are computed over 1000 noise realisations.

Figure 3(c) shows that whilst the NI is generally positive for the additive coupling, it is usually negative for the diffusive coupling. This implies that node activities drive seizures in the network if the coupling is additive, but tend to prevent seizures if the coupling is diffusive. The higher the number of connections of the node, the stronger its ability of driving (preventing) seizures if the coupling is additive (diffusive). We note that as expected for symmetric networks, where the removal of one node is topologically equivalent to the removal of one of the two other nodes, the NI for each node is the same, i.e., within error bars (see e.g. network 8). Overall, we note that the NI distributions are very different, in some cases opposite, for each type of coupling. Moreover, we show the node with the highest NI is different for each network depending on the coupling function.

### Networks

We now turn our attention to larger networks. In this section we compare the role of the different coupling functions on the transient dynamics of the bi-stable model in networks with 64 nodes using the concepts of BNI and NI. To aid our interpretation of the following numerical results we note the relationship between the coupling function and node degree. The mixed coupling function (5) can be rewritten as

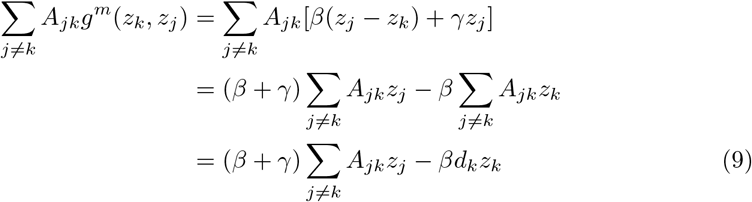

where *d_k_* is the in-degree of node *k*. For simplicity of interpretation, we assume that *z_k_*(*t*) is a positive variable, such as the amplitude *R_k_* (or as the node output in the theta model [14] described in the supplementary material). As above we consider only *β* and *γ* positive. With this set up, the first term in (9) promotes the increase of activity *z_k_*, whereas the second term may only suppress it. Therefore, the additive coupling models excitation, whereas the diffusive coupling may model both excitation or inhibition depending on node activities and network structure.

Substituting (9) into (1) gives

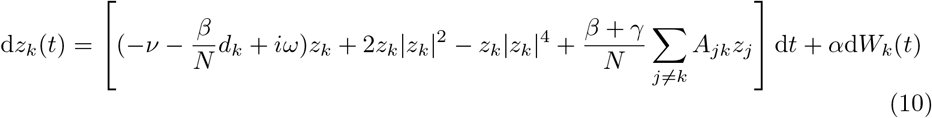

This shows that inhibition is node dependent, being proportional to a node’s own activity and number of in-connections. This effect will lead to very different node behaviour in networks where the in-degree is highly heterogeneous between nodes.

The difference between additive and diffusive coupling is particularly distinct in all-to-all networks (*A_jk_* = 1 for all *j* ≠ *k*). In these networks, all nodes are topologically equivalent with in-degree *d_k_* = (*N* – 1), and if the network is sufficiently large, then their activity is on average the same. As a consequence, before any node escapes, the diffusive term, 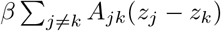, is approximately zero. In contrast, the additive term, 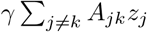 is approximately equal to *γ*(*N* – 1)*z_k_*. This suggests that the diffusive coupling tends to have a weak influence on the dynamics of resting well-connected networks, whereas the additive coupling term tends to be stronger as the number of connections increase.

Below we consider large random and scale-free networks, both directed and undirected. Whilst random networks are fairly homogeneous with regards to the degree distribution, scale-free networks are highly heterogeneous, having some highly connected nodes [26].

#### Brain network ictogenicity

We first focus on the BNI, the network’s propensity to generate seizure activity, across different coupling functions and network structures. Figure 4 shows the BNI as a function of the coupling strength *γ* when using the additive coupling (4). We chose a range of *γ* such that we could observe the greatest overall possible variation in BNI, and considered five levels of noise *α* to show its impact on BNI. We observe that in all considered network structures, the BNI grows monotonically with *γ*. This result means that in all these types of networks, the stronger the connection strength between nodes, the more likely the network is to generate seizure activity. This result is in agreement with our observations in the two-nodes motifs where the first and second escape times decrease as *γ* increases (see Fig. 2), as well as the BNI results in the three-node networks (see Fig. 3). We also observe that the higher *α* is, the higher the BNI is. Both the coupling and noise terms are positive, and at larger values the nodes are more likely to transit to the seizure state, hence increasing the BNI. The BNI dependence with *γ* is qualitatively similar across the four types of network topologies, although we note that in directed scale-free networks the growth in BNI is not as steep as in the other networks and there is greater variability across network realisations. This observation is presumably a consequence of directed scale-free networks having the most heterogeneous degree distributions across all networks considered. The BNI appears to plateau at values lower than 1 because there may be nodes that are unreachable by the influence of their neighbours (i.e., nodes with only outgoing connections), which remain in the resting state. S1 Fig shows that the results are qualitatively similar in networks with mean degree of 8. The main difference in the BNI curves of the networks with higher mean degree is that they are steeper than those of networks with lower mean degree. This result is to be expected: higher mean degree implies on average stronger influence from the coupling term and so the BNI reaches it maximum at lower values of *γ*.

**Fig 4.**
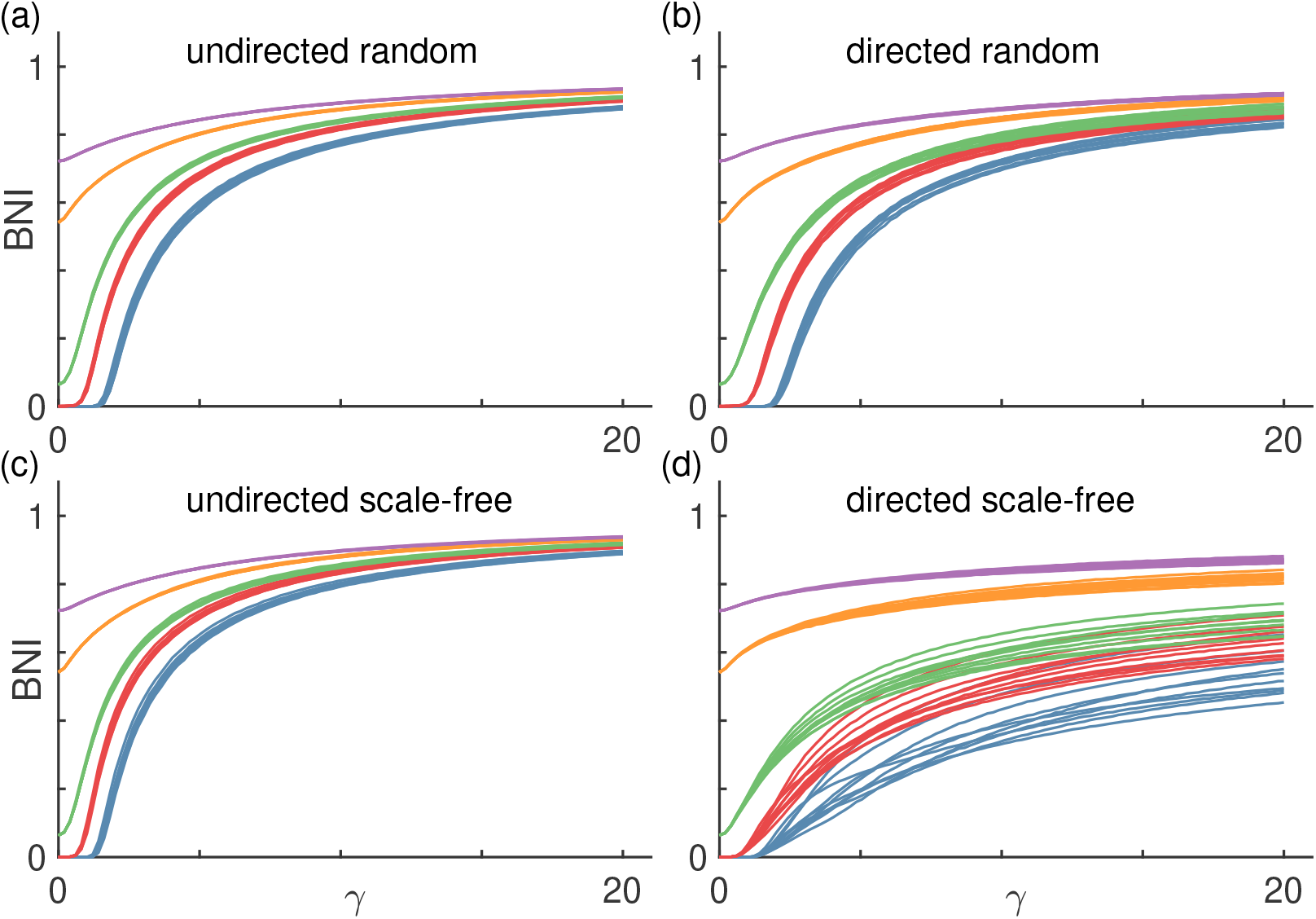
BNI as a function of additive coupling strength *γ*. Each panel shows BNI curves for different network topologies: (a) undirected random networks, (b) directed random networks, (c) undirected scale-free networks, and (d) directed scale-free networks. Each color corresponds to a different level of noise *α*: blue is *α* = 0.001, red is *α* = 0.005, green is *α* = 0.01, orange is *α* = 0.03, and purple is *α* = 0.05. Finally, each curve corresponds to a different network realisation. We used 10 network realisations per network topology and the mean degree of all networks is *c* = 4; see Methods for details.

Figure 5 shows the BNI as a function of the coupling strength *β* when using the diffusive coupling (3). We observe striking differences relative to Fig. 4 depending on the network type as well as coupling strength. As in the case of the additive coupling, we chose a range of *β* such that we could observe a full variation in BNI, and chose five values of the noise amplitude *α*. For *α* = 1 we note that the dynamics of each network are noise dominated leading to a BNI of 1, where all nodes transition to the seizure state within the simulation time, for all values of *β*. First, in the case of undirected networks, we find that the BNI decreases monotonically with *β*. The increase of the coupling strength promotes the inhibitory effect of each node and decreases the BNI. Higher values of noise amplitude *α* imply that the BNI is higher at *β* = 0. On one hand, we had to use higher *α* values for simulations with diffusive coupling relative to those with additive coupling such that we could observe BNI> 0. On the other hand, we used a range of values for *β* four times higher than the range for *γ*, because while the additive coupling cooperated with the noise in driving excitation, the diffusive coupling opposed the noise to suppress activity, thus requiring higher noise magnitude. Second, directed networks are characterised by BNI curves that are not always monotonically decreasing. All directed scale-free and some directed random networks show a local minimum in the BNI curve at low *β* values, followed by a local maximum and a plateau (for the examples where *α* < 0.1). This illustrates how the diffusive coupling may model both excitation or inhibition depending on network structure, whereas the influence of the additive coupling is only excitatory. Third, we observe higher variability in the BNI curves across undirected network realisations with diffusing coupling compared to additive coupling. Also, we observe considerable variability in the BNI curves across the directed random network realisations, and little variability across the directed scale-free network realisations, which is the opposite relation in terms of variability observed for the additive coupling. However, there is one similarity in terms of the BNI curves of the directed scale-free networks between additive and diffusive coupling: both sets of curves appear to plateau at some intermediate BNI value, presumably due to the existence of ’unreachable’ nodes. S2 Fig shows that the results are similar in networks with higher mean degree. As observed with the additive coupling, the BNI curves are steeper. These results suggest that increasing the mean degree is effectively similar to increasing the coupling strength (regardless of the type of coupling).

**Fig 5.**
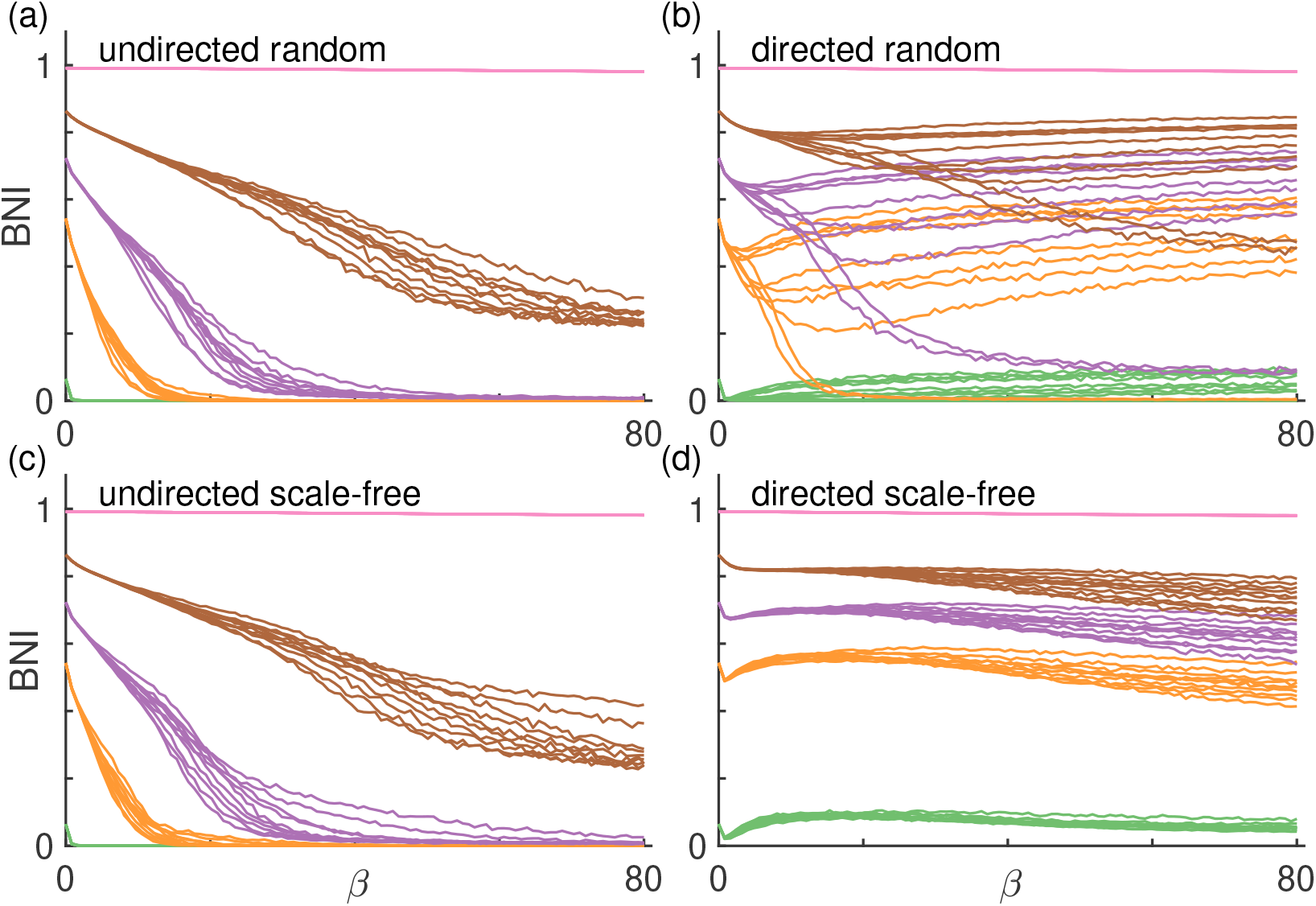
BNI as a function of diffusive coupling strength *β*. Each color corresponds to a different level of noise *α*: green is *α* = 0.01, orange is *α* = 0.03, purple is *α* = 0.05, brown is *α* = 0.1, and pink is *α* = 1. All other parameters are the same as in Fig. 4.

Figure 6 shows the BNI as a function of the coupling strength *γ* (with fixed (*β*) and *β* (with fixed *γ*) when using the mixed coupling with each of the four network types. As in the additive coupling case, the BNI grows monotonically with increasing *γ*, and, as in some of the diffusive coupling cases, the BNI decreases monotonically with increasing *β* in all of the networks considered. These curves can be considered as cross sections of a generalised BNI surface in a *γ* – *β* plane (or a hypersurface if we consider a range of *α* values). Such surface would contain the curves observed in Figs. 4, 5, and 6 for a given network and a fixed *α*. Interestingly, Fig. 6(b) suggests that a sufficiently strong additive component *γ* can prevent the diffusive coupling component of suppressing the BNI. Note that at *α* = 0.05, the diffusive coupling was capable of reducing the BNI to zero in the case of the undirected networks, see the purple curves in Figs. 5(a) and (c). We also observe higher agreement between the BNI curves across different network topologies with mixed coupling than with diffusive coupling alone. In other words, this result suggests that the additive coupling component causes the BNI curves to be more uniform across network topologies relative to the diffusive coupling component of the mixed coupling.

**Fig 6.**
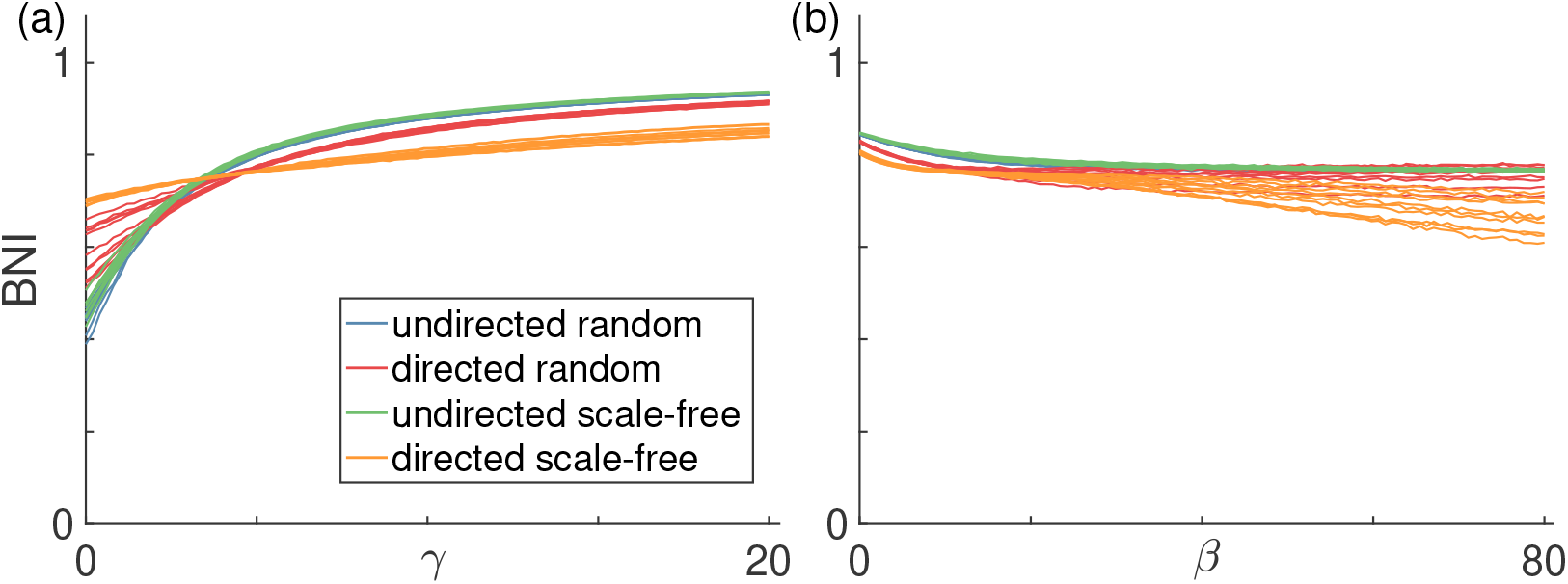
BNI as a function of (a) *γ*(*β* = 10) and (b) *β*(*γ* = 5) using the mixed coupling. Each color corresponds to a different network topology: blue corresponds to undirected random networks, red to directed random networks, green to undirected scale-free networks (overlaps the blue), and orange to directed scale-free networks. Each curve corresponds to a different network realisation. We fix *α* = 0.05 and mean degree *c* = 4.

#### Node ictogenicity

The NI quantifies the contribution of each node to the overall network’s propensity to generate seizures. Identifying the nodes with the highest contribution (i.e., highest NI) can be useful to inform epilepsy surgery [6, 14, 34]. Figure 7 shows representative NI distributions for each of the four network topologies, for a given level of noise. These representative NI distributions show how the NI values depend on the coupling function.

**Fig 7.**
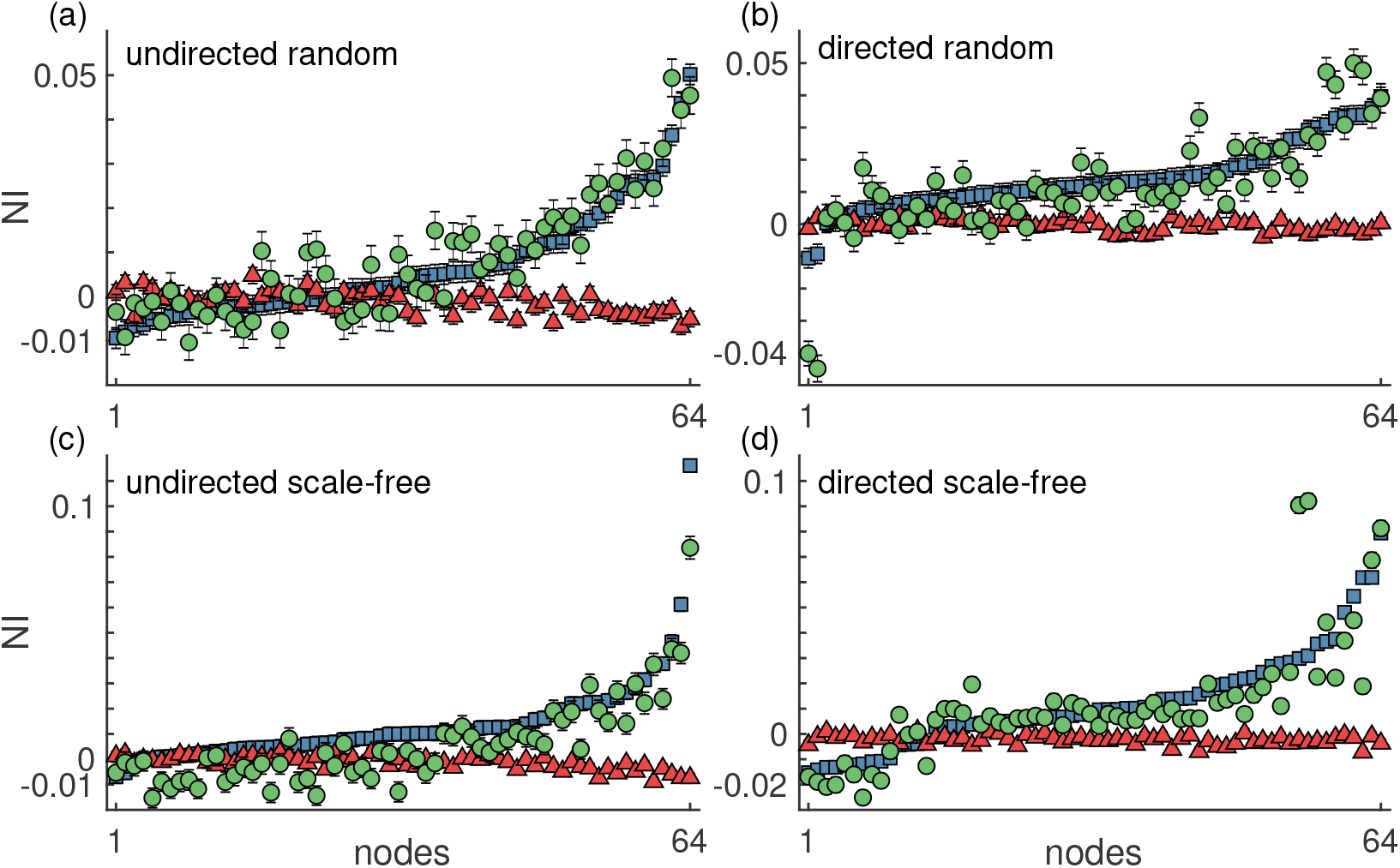
Representative NI distributions of (a) undirected random, (b) directed random, (c) undirected scale-free, and (d) directed scale-free networks using the different couplings. The blue squares represent the NI computed using the additive coupling, the red triangles correspond to the diffusive coupling and the green circles correspond to the mixed coupling. The nodes were sorted such that the NI grows monotonically for the additive coupling. The error bars represent the standard error across 1000 realisations. We used *α* = 0.005 for the additive coupling, *α* = 0.03 for the diffusive coupling, and *α* = 0.01 for the mixed coupling. In all three coupling cases, parameters *γ* and *β* were chosen such that BNI_pre_ = 0.5; for the mixed coupling, we fixed *β* = 10 and chose *γ*. All networks had mean degree *c* = 4.

We observe that the NI when using the additive coupling has a much larger range, NI∈ [–0.01,0.1], over the nodes than the range, NI∈ [–0.01, 0.01], for the diffusive coupling. Furthermore, the absolute values of NI are generally higher in the additive case relative to the diffusive case. A large proportion of the nodes have a positive NI with the additive coupling, meaning that the removal of nodes contributes to an overall reduction in BNI, whereas the majority of nodes with diffusive coupling have negative NI values, which implies that removing nodes can increase the network’s ability to generate seizure activity. We find that for the diffusive coupling nodes with the highest degree are more likely to have the lowest NI. From (9) we observe that inhibition is node dependent, being proportional to a node’s own activity and number of in-connections. In contrast, the NI is highest in highly connected nodes when using the additive coupling because such nodes are more likely to be both excited into the seizure state and to be capable of exciting their neighbours.

Figure 7 also shows the NI distribution computed with the mixed coupling, which in each panel follows the NI distribution from the additive coupling, suggesting that the additive component of the mixed coupling is dominant for the chosen parameters. We found consistent results using other network realisations and other *β* and *γ* parameter values.

To better compare NI distributions from the different coupling functions, we computed the weighted Kendall correlation rank *τ*. Figure 8 shows that the NI distributions from additive and diffusive couplings are not just different, they actually tend to rank the nodes in opposite order. Nodes with the highest NI in the additive coupling are likely to be the nodes with the lowest NI in the diffusive coupling, and vice versa, as observed in the three-node networks (see Fig. 3(c)). Additionally, as observed in Fig. 7, the NI orderings of the additive and mixed couplings are in almost perfect agreement (average *τ* > 0.96 in all types of networks) for the chosen parameters. Consequently, the relationship between the diffusive and mixed couplings is similar to the additive and mixed couplings as assessed by *τ*. We expect that as the diffusive component of the mixed coupling would be increased (and/or the additive component would be decreased), the *τ* value relating the diffusive and mixed couplings would increase, and the *τ* value comparing the additive and mixed couplings would decrease. These results are consistent across all network topologies, with lower *τ* values in the undirected networks relative to directed networks when comparing the diffusive coupling to the other couplings. S3 Fig, S4 Fig, and S5 Fig complement these findings by showing that the NI is related to the number of connections that each node has. We find a positive correlation between NI and node degree in the additive and mixed couplings (see S3 Fig and S5 Fig), whereas the correlation is negative in the diffusive coupling (see S4 Fig). Thus, nodes with higher degree have higher NI in the additive coupling, but lower NI in the diffusive coupling case. The mixed coupling can presumably range between the two extremes depending on parameters.

**Fig 8.**
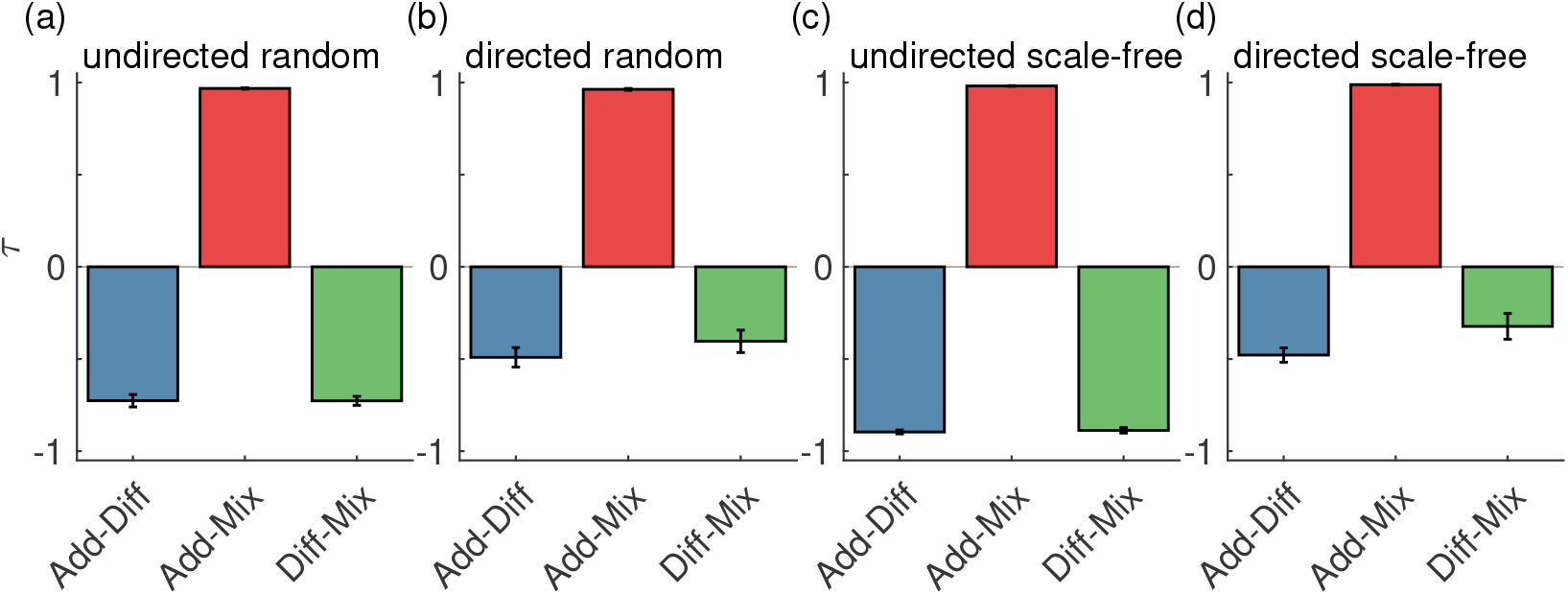
Comparison of NI distributions using different couplings. The weighted Kendall correlation rank *τ* quantifies the consistency of ordering nodes according to their NI values when using the different couplings. The blue bars correspond to the comparison between additive and diffusive coupling, the red bars correspond to additive versus mixed coupling, and the green bars to the diffusive versus mixed coupling. Different panels show the comparison for different network topologies: (a) undirected random, (b) directed random, (c) undirected scale-free random, and (d) directed scale-free networks. The error bars represent the standard error across the 10 network realisations per network topology. All parameters are the same as in Fig. 7.

### BNI of MEG functional networks

To further assess the impact of choosing either additive or diffusive coupling in studies that aim to investigate the emergence of seizures on real-world brain networks, we computed the BNI of 26 MEG functional networks from people with JME and 26 from healthy controls using both additive and diffusive coupling. Figure 9 shows that the BNI based on additive and diffusive couplings rank the individuals in reverse order. Individuals with the highest ’additive BNI’ have the lowest ’diffusive BNI’, and vice versa. Furthermore, while most individuals have similar diffusive BNI, they are well distinguished in terms of additive BNI. Finally, we observe that people with JME have on average higher additive BNI than controls (Mann–Whitney U test, *p* = 0.0062), but lower diffusive BNI than controls (*p* = 0.0026). It is important to note that higher BNI in people with JME relative to controls is the expected, given that higher BNI is assumed to characterize a brain network with a higher propensity to produce seizures. Therefore, only the additive coupling provides the hypothesized BNI distinction between the two groups. We performed the same comparison for other *γ* and *β* values and found similar results (not shown).

**Fig 9.**
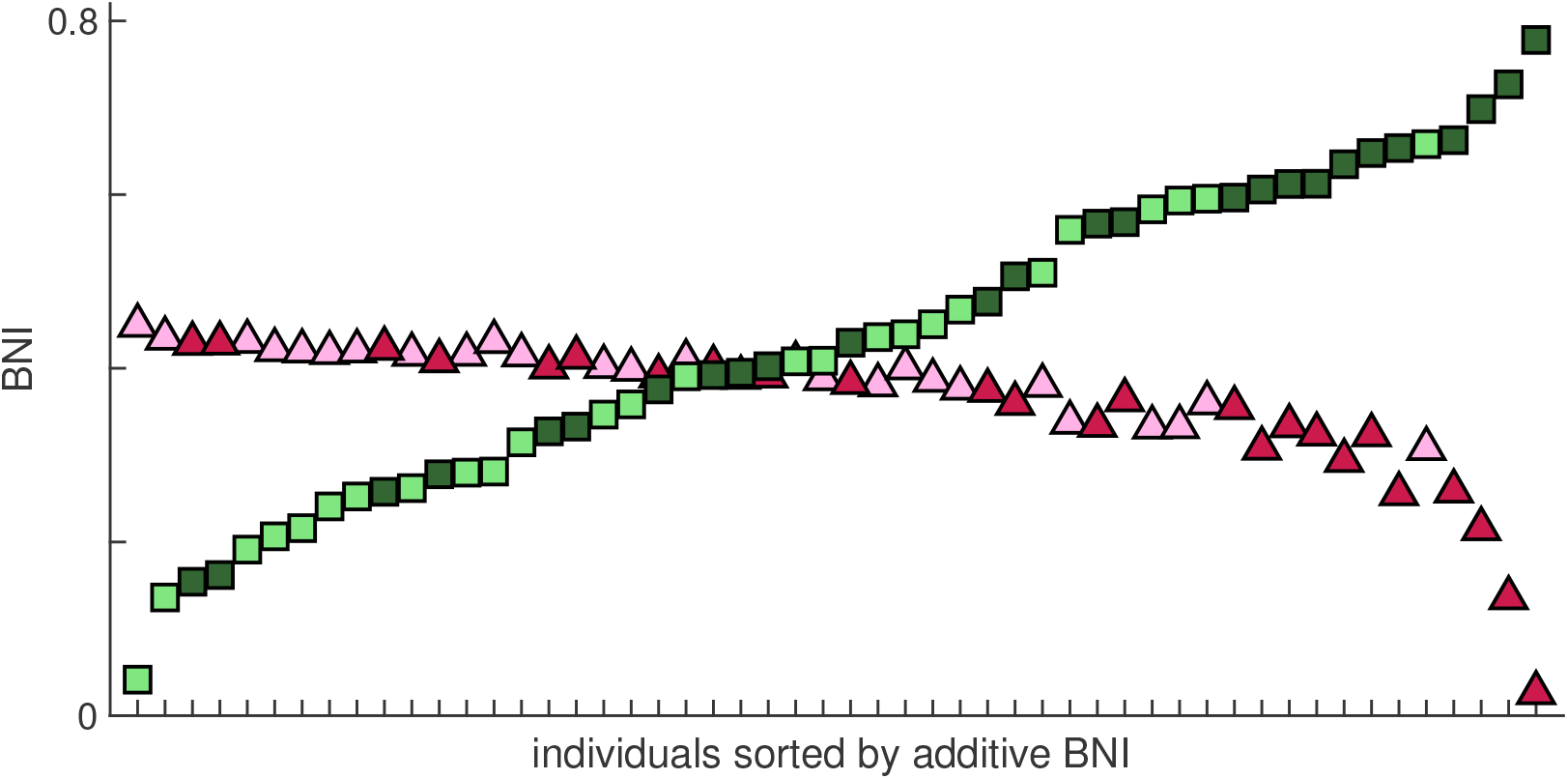
Comparison of BNI using additive and diffusive couplings on MEG functional networks. Each square (triangle) is the BNI value based on additive (diffusive) coupling of each individual. The individuals were sorted such that the BNI values based on the additive coupling were monotonically increasing. Light green squares and light pink triangles correspond to healthy controls, and dark green squares and dark pink triangles correspond to people with JME. We used *α* = 0.005 and *γ* = 15 for the additive coupling simulations, and *α* = 0.03 and *β* = 4.6 for the diffusive coupling simulations.

## Discussion

BNMs are useful tools to understand the role of brain network structure on healthy and pathological brain dynamics. These models make assumptions about how brain regions behave and how they interact. Here, we investigated two main modelling choices of interaction, the additive coupling and the diffusive coupling, aiming to understand their role on simulated brain dynamics. We focused on a bi-stable model of seizure transitions, a model that has been used in the literature with both additive and diffusive couplings [11–13, 16, 17]. We further considered a mixed coupling, combining the additive and diffusive couplings. We analysed the model on artificial networks with two, three and 64 nodes, as well as MEG functional networks, and characterised the dynamics using escape time, BNI and NI. These measures have been used in the literature to make predictions about ictogenicity. We showed that additive and diffusive couplings provide fundamentally different, often contradictory, predictions in both the large artificial and MEG functional networks. The mixed coupling gave results in between the two depending on parameters. For two connected nodes, we demonstrated that additive and diffusive couplings give rise to two different bifurcation diagrams, particularly at large coupling values, which explains the different dynamical behaviours in terms of escape times. In three-node networks, we found that generally networks with additive coupling have higher BNI than networks with diffusive coupling, and the difference in BNI increases in networks with more connections. Furthermore, nodes with higher NI with the additive coupling had the lowest NI with the diffusive coupling. Our simulations in larger networks with 64 nodes further supported these observations. Finally, we applied the BNI to MEG functional networks from people with JME and healthy controls using both additive and diffusive couplings. We found that people with JME had a higher BNI based on the additive coupling than the controls, and we observed the opposite using the diffusive coupling.

Our observations that the BNI and NI are different depending on whether we use additive or diffusive coupling have consequences for studies aiming to apply these measures to interrogate functional networks obtained from clinical data, as illustrated with our BNI results on the MEG networks. The BNI has been used to differentiate the functional networks of healthy people from people with epilepsy inferred from resting-state data [11, 12, 31]. The hypothesis is that people with epilepsy have a higher enduring propensity to generate seizures than healthy individuals, and that this propensity can be assessed from their resting-state functional networks. The BNI is meant to assess this enduring feature of the epileptic brain and, therefore, it is hypothesized that people with epilepsy have a higher BNI than healthy controls.However, our findings based on artificial networks suggest that whether the BNI is higher in one group relative to the other depends on the choice of coupling function. Group A may have higher BNI than group B if we choose the additive coupling, whereas group B may have higher BNI than group A if we choose the diffusive coupling. We confirmed this expectation by computing the BNI on the MEG functional networks using the two couplings. We observed that the BNI using the diffusive coupling provided almost the opposite BNI ranking of individuals compared to the BNI using the additive coupling. In particular, we found that only the BNI based on the additive coupling distinguished the JME group with higher BNI values than the healthy group. Together with previous evidence [12, 31], this result suggests that the additive coupling may be a better modeling choice than the diffusive coupling, for the purpose characterizing ictogenic brain networks with BNI.

The NI has been used to model epilepsy surgery and to make predictions about the epileptogenic zone [6, 14, 16, 17, 34]. Nodes with the highest NI are taken as predictors of the epileptogenic zone. The fact that the NI distribution strongly depends on the coupling choice implies that predictions about the epileptogenic zone also depend on this choice. The nodes with the highest NI in the additive coupling are likely to be the nodes with the lowest NI in the diffusive coupling. Such disagreement between additive and diffusive couplings highlights the need of finding which coupling choice is most appropriate to model the brain’s ictogenicity. Future work may attempt to answer this question by fitting the model with the mixed coupling function to electrophysiological recordings using a search over the full (*γ*, *β*) parameter space. A more detailed study may even consider weighted networks in which different connections are characterised by different *γ* and *β* values. However, we note that most studies that have used the bi-stable model or other models of ictogenicity to investigate data have used the additive coupling [6, 12–14, 34]. Their promising results suggest that the additive coupling, or perhaps the mixed coupling, may be more appropriate than the diffusive coupling for such investigations.

We highlight that our simulations based on the additive and diffusive couplings provide different understandings about the role of single node dynamics and network structure on ictogenicity. In the case of the additive coupling, ictogenicity can result from pathological single nodes and/or pathological brain structure. In this context, ’pathological single nodes’ are nodes that are able to generate seizure activity even in isolation. Within the phenomenological model framework, such nodes have been modelled by making either *v* or *α* node specific [16, 21]. With additive coupling, pathological single nodes may cause seizures due to the excitatory nature of all interactions. However, pathological single nodes are not necessary for a network to generate seizures. The network ictogenicity may result from the network’s structure. For example, highly connected nodes are likely to be prone to generate seizures, which in turn makes the whole network prone to seizures. Given the excitatory nature of the additive coupling, the network structure can only enhance the ictogenicity of individual nodes. In contrast, in the case of diffusive coupling, ictogenicity is the result of both pathological nodes and pathological brain structure. A network without pathological nodes, i.e., nodes capable of generating seizures in isolation, cannot generate seizures. In our simulations, we had to choose a sufficiently large level of noise *α* so that the BNI curves had a non-zero BNI at *β* = 0, otherwise, the BNI would be zero at all values of *β*. On the other hand, the existence of pathological nodes does not guarantee network ictogenicity because certain network structures may prevent ictogenicity due to the potential inhibitory role of the diffusive coupling. Thus, seizures may only emerge in networks with diffusive coupling if the network contains pathological nodes and if the network structure is such that enables their pathological activity. As a consequence, our results suggest that if the additive coupling is a better model of large-scale brain interactions, then knowledge about network structure may be sufficient to assess brain ictogenicity because pathological nodes are not necessarily required to drive seizures. On the other hand, if the diffusive coupling is a better approximation of large-scale brain interactions relevant for epilepsy, then knowledge about network structure is insufficient to assess brain ictogenicity. Unfortunately, evaluating whether single nodes are pathological remains a challenge.

The results presented in the main text using the bi-stable model of seizure transitions are supported by our results presented in the Supplementary Material using the theta model [14]. The theta model is an alternative model of ictogenicity and we used it to test whether our comparison between additive and diffusive couplings was model dependent. The findings presented in the Supplementary Material show that the two models are in agreement. Not only the relation between results obtained with additive and diffusive couplings is similar, but also the two models provide similar results when using the same coupling functions (see S12 Fig). These results are in agreement with previous studies comparing these and other models using the additive coupling [20]. Also, the relation between ictogenicity and the number of connections uncovered in our analysis is also in agreement with previous findings using both additive [14] and diffusive coupling [17].

We emphasise that our results should be broadly relevant for studies using BNMs beyond their application to epilepsy. The fact that the coupling choice crucially defines the escape time, BNI and NI suggests that other measures of network dynamics may also be affected. For example, Hansen *et al*. [5] used a BNM with additive coupling to simulate functional connectivity dynamics on structural brain connectivity. Demirtaş *et al*. [7] used a BNM with diffusive coupling to study the mechanisms responsible for connectivity changes in Alzheimer’s disease. Also, Cabral *et al*. [35] used a BNM based on Kuramoto oscillators with diffusive coupling to investigate the emergence of resting-state functional connectivity. All these and other studies’ conclusions may be questionable given their likely dependence on the coupling choice. Furthermore, our results have potentially wide-reaching implications for studies that aim to establish biophysical models of large-scale brain interactions [36].

## Conclusion

Here we compared the impact of using additive or diffusive coupling on the dynamics of a BNM relevant for epilepsy. We showed that the two couplings are not interchangeable. On the contrary, the dynamics on the networks are different and the predictions of node and network relative ictogenicity are often opposite. We used the two coupling frameworks to assess resting-state functional networks inferred from MEG from people with JME and healthy controls and found opposing results in terms of network’s propensity to generate seizures. The additive coupling provided the hypothesized result of higher ictogenic propensity on brain networks from people with JME relative to networks from controls. Thus, our results and evidence from the literature suggest that the additive coupling may be a better modeling choice than the diffusive coupling, at least for BNI and NI studies. Future BNM studies should motivate and validate the choice of coupling to properly model brain activity and to obtain reliable predictions about brain function and dysfunction.

## Supporting information

**S1 Fig.**
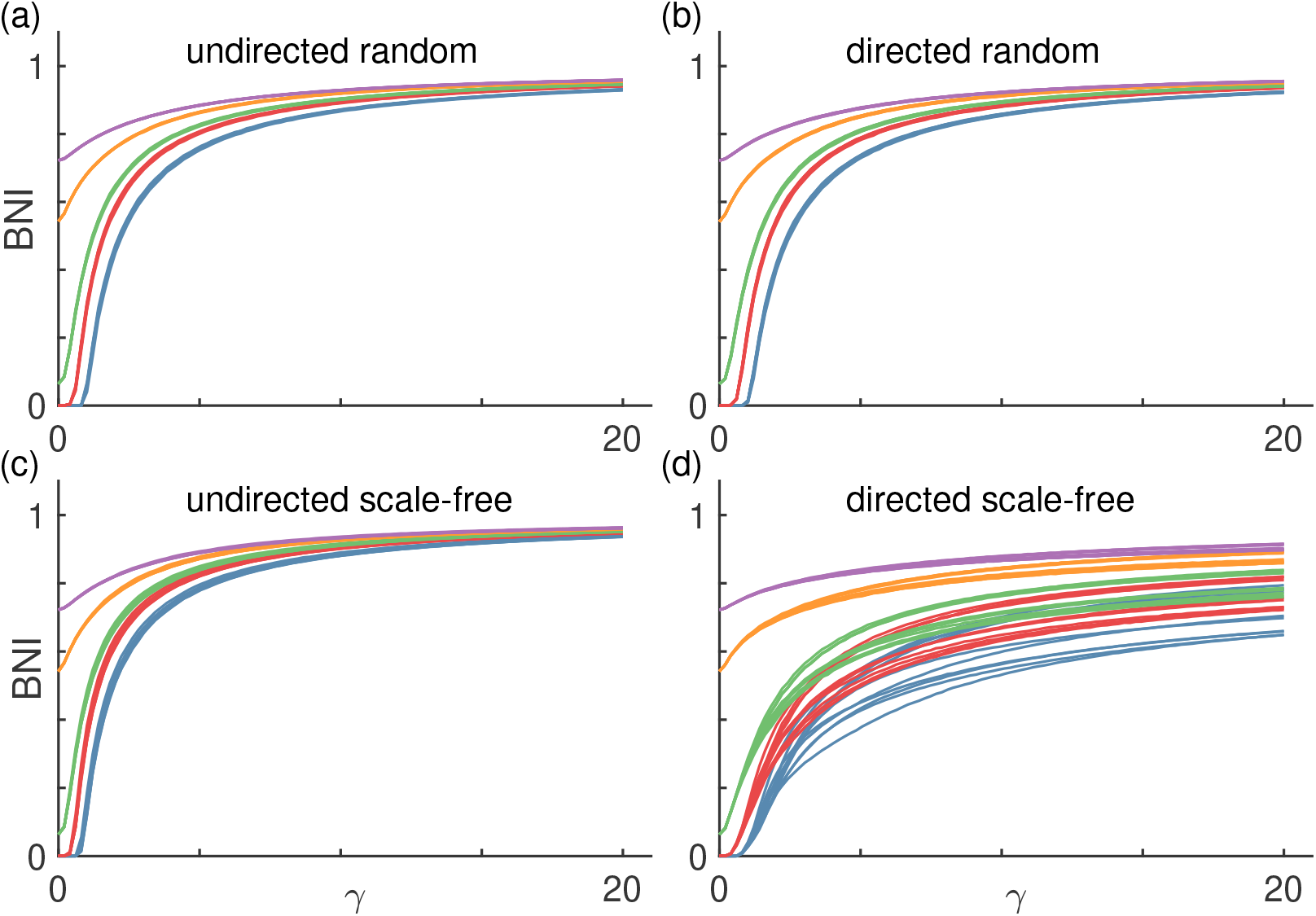
BNI as a function of *γ* using the additive coupling. As in Fig. 4, each panel shows BNI curves for different network topologies: (a) undirected random networks, (b) directed random networks, (c) undirected scale-free networks, and (d) directed scale-free networks. Each color corresponds to a different level of noise *α*: blue is *α* = 0.001, red is *α* = 0.005, green is *α* = 0.01, orange is *α* = 0.03, and purple is *α* = 0.05. Finally, each curve corresponds to a different network realisation. We used 10 network realisations per network topology. The mean degree of all networks is *c* = 8.

**S2 Fig.**
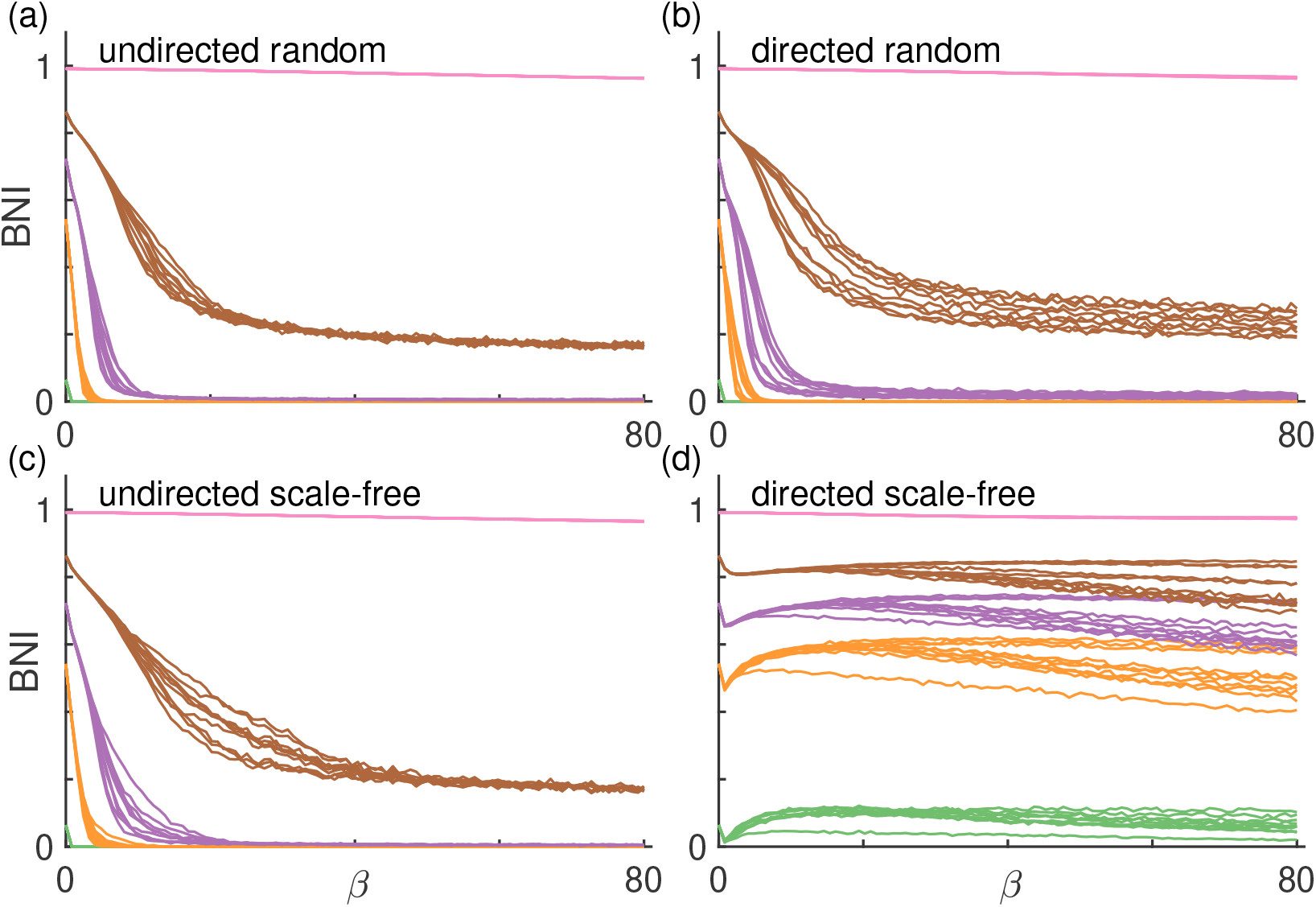
BNI as a function of *β* using the diffusive coupling. As in Fig. 5, each panel shows BNI curves for different network topologies: (a) undirected random networks, (b) directed random networks, (c) undirected scale-free networks, and (d) directed scale-free networks. Each color corresponds to a different level of noise *α*: green is *α* = 0.01, orange is *α* = 0.03, purple is *α* = 0.05, brown is *α* = 0.1, and pink is *α* = 1. Finally, each curve corresponds to a different network realisation. We used 10 network realisations per network topology. The mean degree of all networks is *c* = 8.

**S3 Fig.**
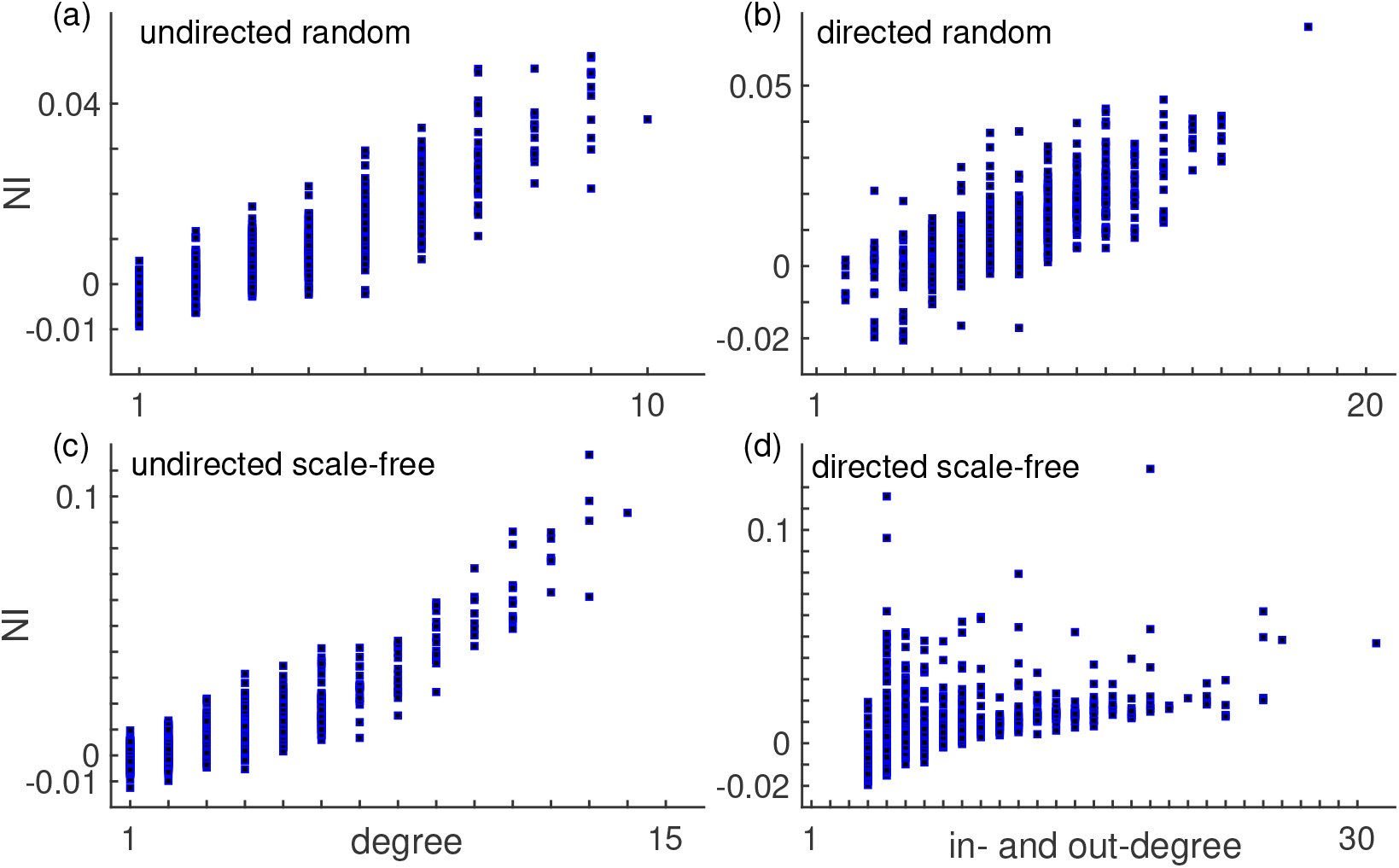
NI versus degree across different network topologies using the additive coupling. Each panel corresponds to a different network topology: (a) undirected random, (b) directed random, (c) undirected scale-free, and (d) directed scale-free networks. In the case of the directed networks, the horizontal axis shows the sum of in- and out-degree. This figure combines the NI distributions across 10 different network realisations per network topology. The average correlation between NI and degree is 0.75 and the average p-value is 0.0015. All parameters are the same as in Fig. 7.

**S4 Fig.**
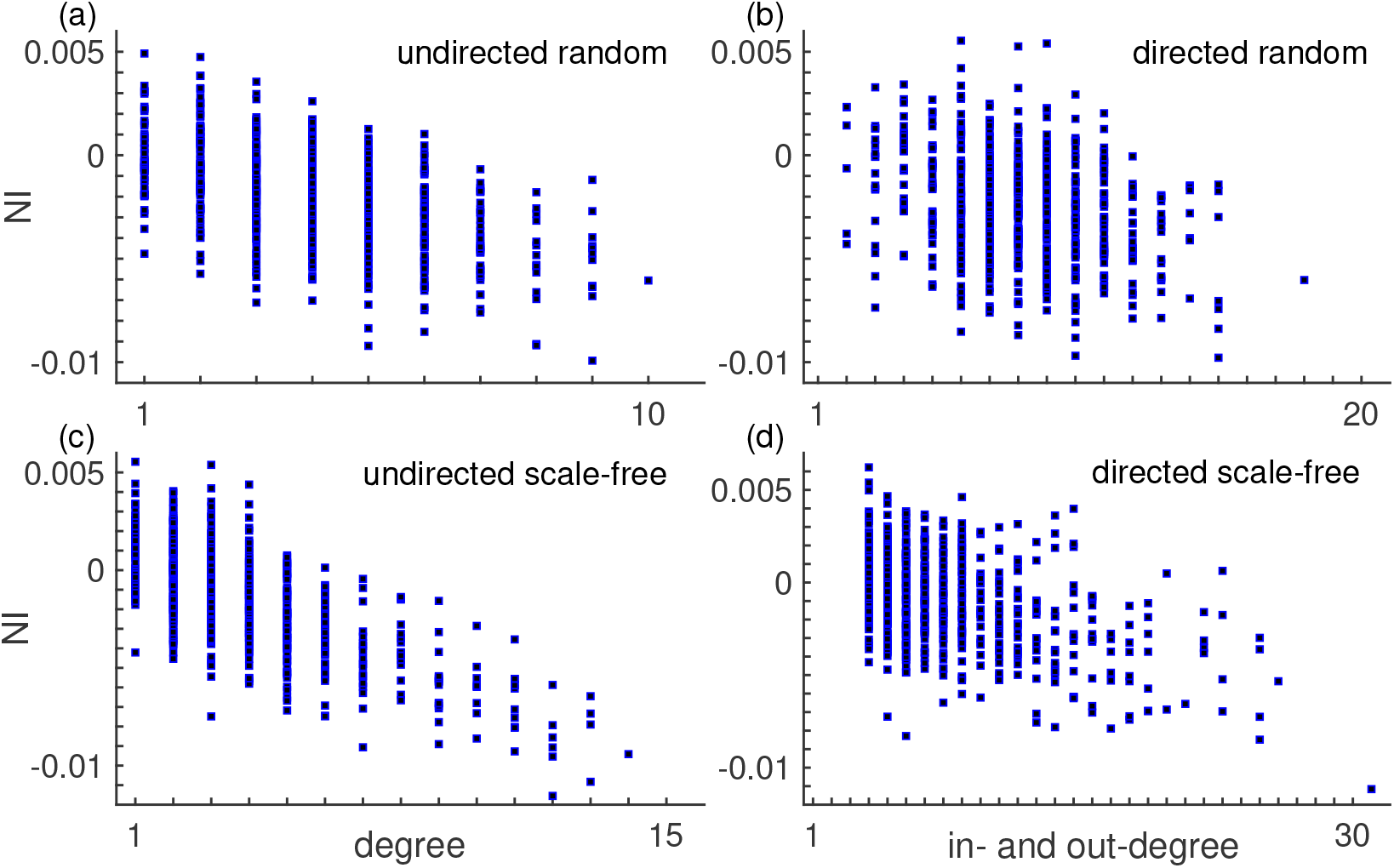
NI versus degree across different network topologies using the diffusive coupling. Each panel corresponds to a different network topology: (a) undirected random, (b) directed random, (c) undirected scale-free, and (d) directed scale-free networks. In the case of the directed networks, the horizontal axis shows the sum of in- and out-degree. This figure combines the NI distributions across 10 different network realisations per network topology. The average correlation between NI and degree is –0.57 and the average p-value is 0.0017. All parameters are the same as in Fig. 7.

**S5 Fig.**
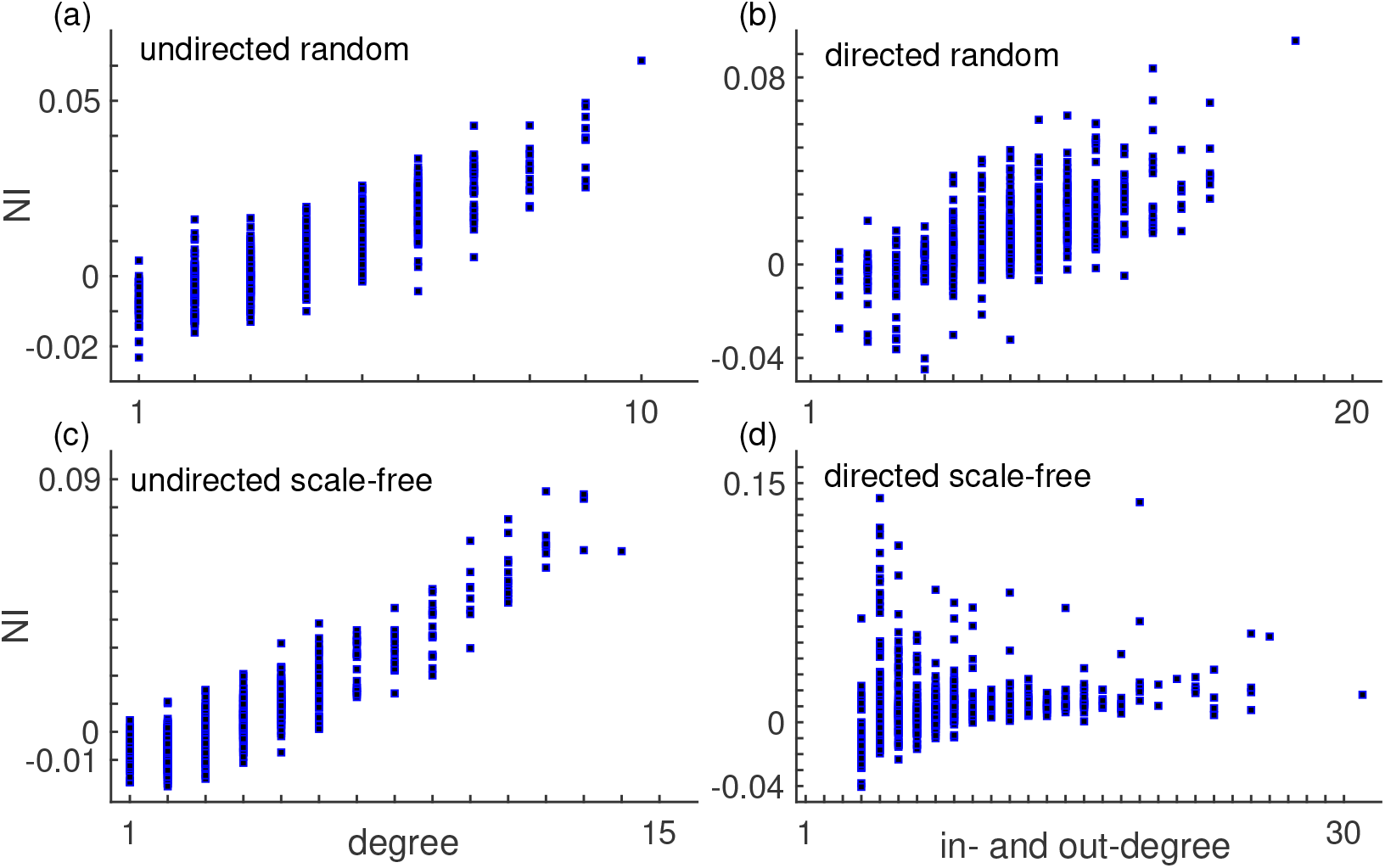
NI versus degree across different network topologies using the mixed coupling. Each panel corresponds to a different network topology: (a) undirected random, (b) directed random, (c) undirected scale-free, and (d) directed scale-free networks. In the case of the directed networks, the horizontal axis shows the sum of in- and out-degree. This figure combines the NI distributions across 10 different network realisations per network topology. The average correlation between NI and degree is 0.69 and the average p-value is 0.040. All parameters are the same as in Fig. 7.

## S1 Appendix. Theta model

In the main text we compare additive, diffusive and mixed couplings within the bi-stable model presented in the Methods. Here, we expand our comparison to a different model of ictogenicity, the theta model [14, 23, 31].

The theta model can be used as a phenomenological BNM of seizure dynamics [14]. The phase *θ_k_* characterises the activity at node *k* and it is described by the differential equation,

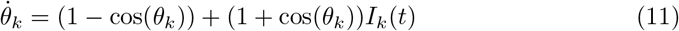

where *I_k_*(*t*) is an input current to the node. At *I_k_* < 0, the node is at a fixed stable phase 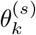 (a ’resting state’). At *I_k_* = 0, there is a saddle-node on invariant circle (SNIC) bifurcation. At *I_k_* > 0, the node oscillates (the ’seizure state’). The current *I_k_*(*t*) is defined by

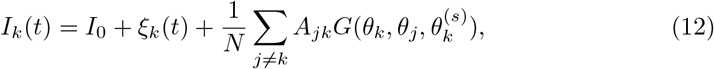

where *I*_0_ is the excitability of node *k*, *ξ_k_*(*t*) are noisy inputs, and the third term on the right hand side is the coupling term. *N* is the number of nodes, *A_jk_* is the adjacency matrix, and 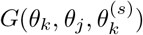 is the coupling function. We consider two coupling functions:

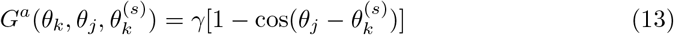

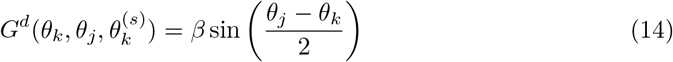

where *G^a^* corresponds to additive coupling, and *G^d^* to diffusive coupling. *G^a^* has been considered in the literature [14, 23, 31, 34], whereas *G^d^* has not.

The phase 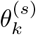 is a stable fixed point obtained from 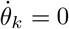 in the absence of noise and node interactions (*ξ*^(*k*)^(*t*) = 0 and *γ* = *β* = 0),

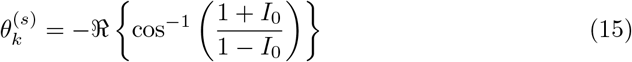

At *I*_0_ > 0 there is no stable fixed point and we use 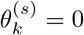.

The noise term is independent across nodes and it is Gaussian distributed with zero mean and variance *σ*^2^. Following previous studies [14, 23, 31, 34], we use *I*_0_ = –1.2 and *σ* = 0.6 in the case of the additive coupling. We consider *σ* = 1.44 in the case of the diffusive coupling. As in the bi-stable model, we had to consider a higher level of noise in the diffusive coupling than in the additive coupling because otherwise the network would always be in the resting state regardless of *β*.

To compare additive and diffusive couplings using this model, we computed the BNI and NI as we did in the main text for the bi-stable model. We also used the same networks (see Section) so that to enable comparisons between the theta and bi-stable models. We integrated the stochastic equations (11) using the Euler-Maruyama method with step size *h* = 10^−2^.

In the theta model, the BNI is defined as follows

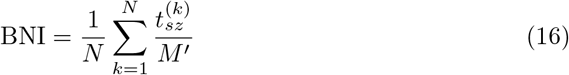

where 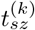 is the time that node *k* spends in the seizure state during a total simulation time *M*′. We used *M*′ = 4 × 10^6^ as in Ref. [23, 31]; see Lopes *et al*. [14] for more details on the calculation of 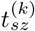. The NI is the same as presented in the main text.

S6 Fig and S7 Fig show the BNI as function of *γ* and *β* for the additive and diffusive couplings respectively. The results are similar to those observed within the bi-stable model. Namely, the BNI grows monotonically with *γ* and decreases with *β*. Also, increasing the mean degree of the networks has a similar effect to increasing *γ* or *β*. A difference between these results and those observed with the bi-stable model is that in the case of the diffusive coupling in directed networks, we found a local minimum in some BNI curves in the bi-stable model (see Figs. 5 and S2 Fig) but not in the theta model. However, we have not performed an exhaustive parameter search for such behaviour in the theta model, therefore we cannot exclude the possibility that it may exist for other parameters.

**S6 Fig.**
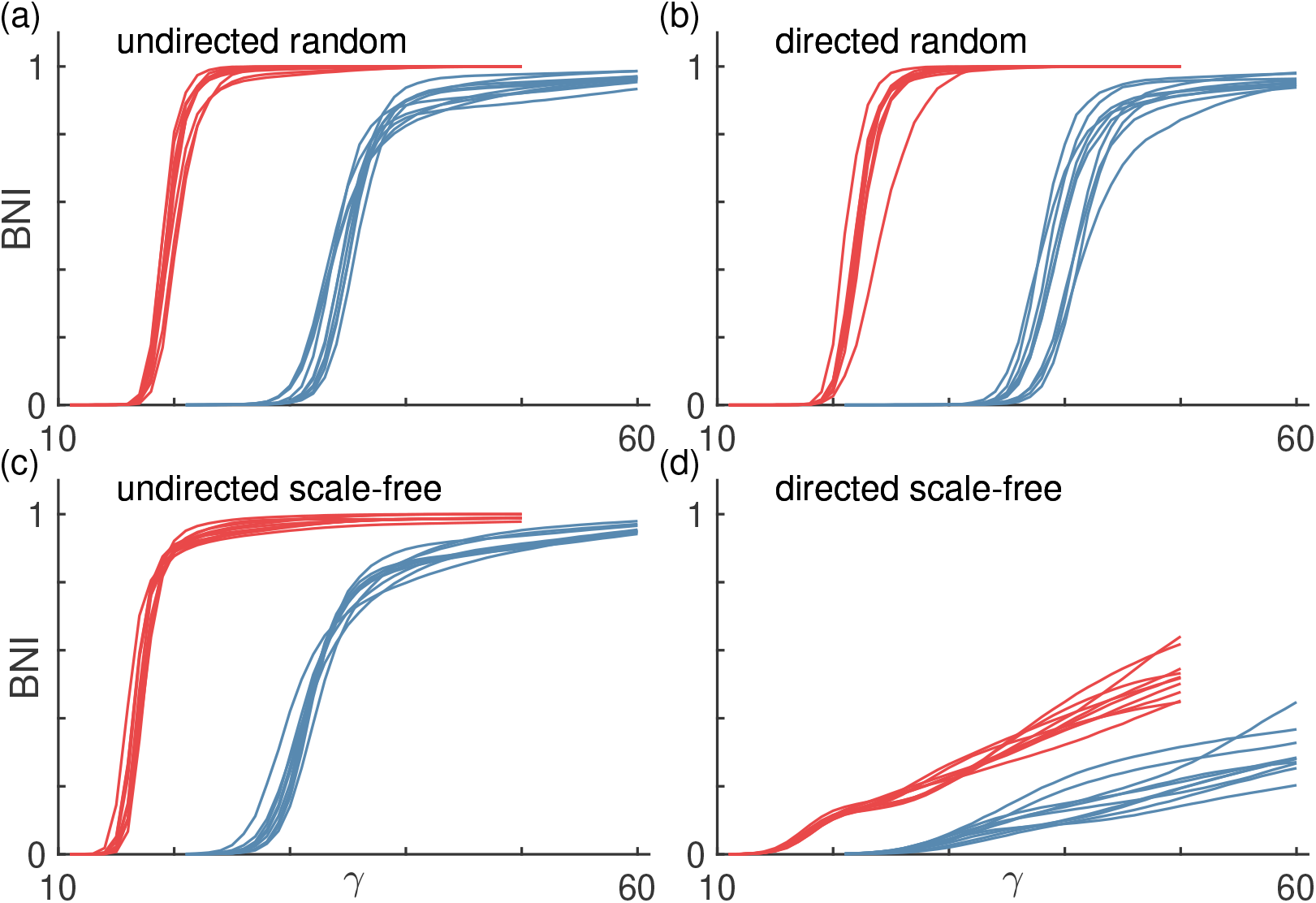
BNI as a function of *γ* using the additive coupling and the theta model. Each panel shows BNI curves for different network topologies: (a) undirected random networks, (b) directed random networks, (c) undirected scale-free networks, and (d) directed scale-free networks. Curves in blue correspond to networks with mean degree *c* = 4, and curves in red correspond to networks with *c* = 8. Each curve corresponds to a different network realisation. We used 10 network realisations per network topology.

**S7 Fig.**
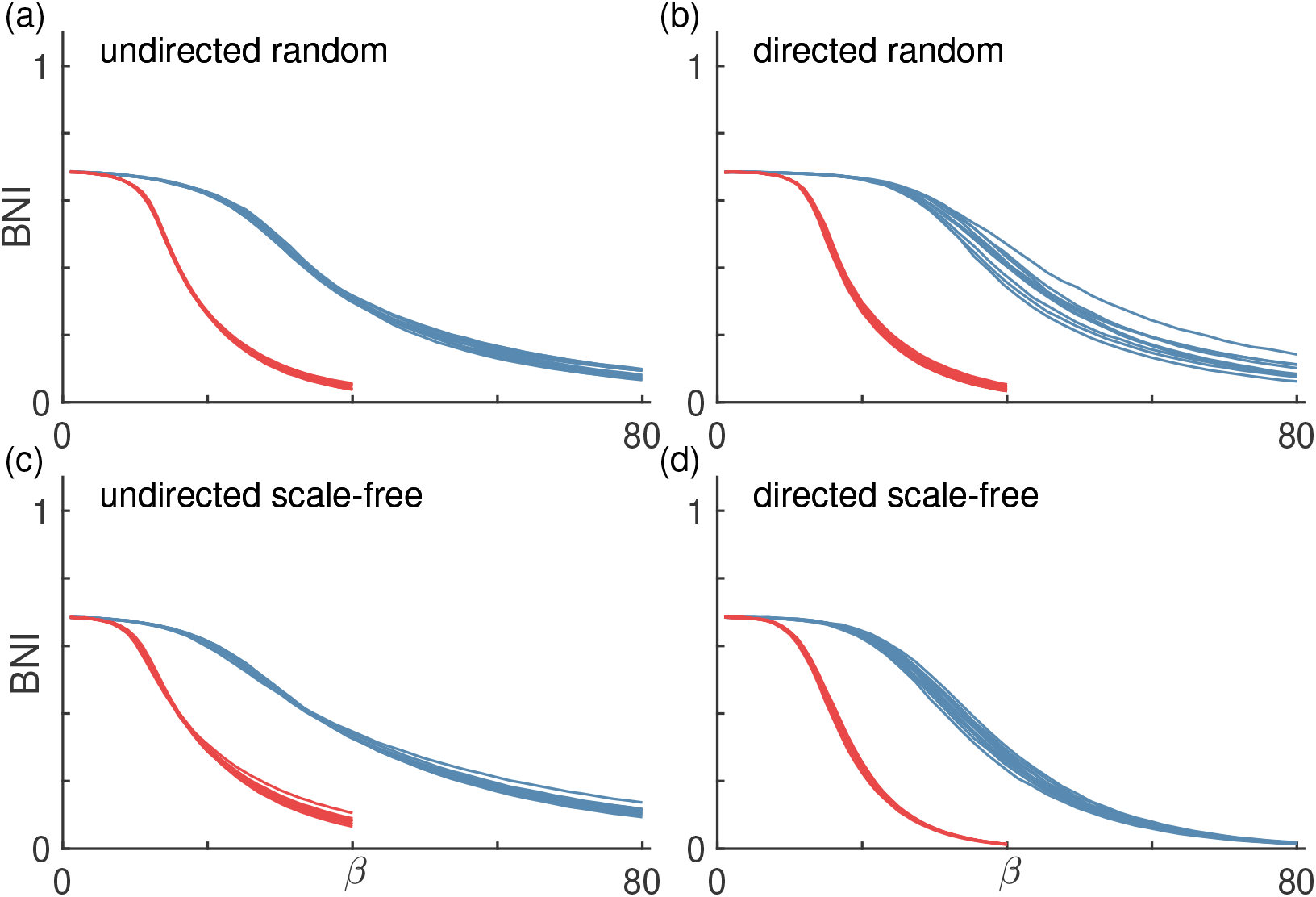
BNI as a function of *β* using the diffusive coupling and the theta model. Each panel shows BNI curves for different network topologies: (a) undirected random networks, (b) directed random networks, (c) undirected scale-free networks, and (d) directed scale-free networks. Curves in blue correspond to networks with mean degree *c* = 4, and curves in red correspond to networks with *c* = 8 (we used a different range of *β* for the two sets of curves). Each curve corresponds to a different network realisation. We used 10 network realisations per network topology.

S8 Fig compares NI distributions computed with additive and diffusive couplings in the theta model. The results are qualitatively the same as those found in the bi-stable model (see Fig. 7). Nodes with higher (positive) NI in the additive coupling case are the nodes with lowest (negative) NI in the diffusive coupling case. Furthermore, the NI in the additive coupling has higher variability than in the diffusive coupling. S9 Fig, S10 Fig, and S11 Fig are also in agreement with the corresponding in the bi-stable model. Briefly, the NI correlates (anti-correlates) with node degree in the additive (diffusive) coupling (except when using additive coupling in directed scale-free networks). The weighted Kendall correlation rank *τ* is close to –1 in all networks except directed scale-free networks, showing that the NI from additive and diffusive couplings rank nodes in reverse order. The directed scale-free networks is presumably an exception due to its highly heterogeneous nature in terms of degree distribution. We speculate that the role of in-degree and out-degree is different in the two coupling cases, but not the ’reverse’, as it seems to be the case when considering undirected networks, where the in-degree is equal to out-degree.

**S8 Fig.**
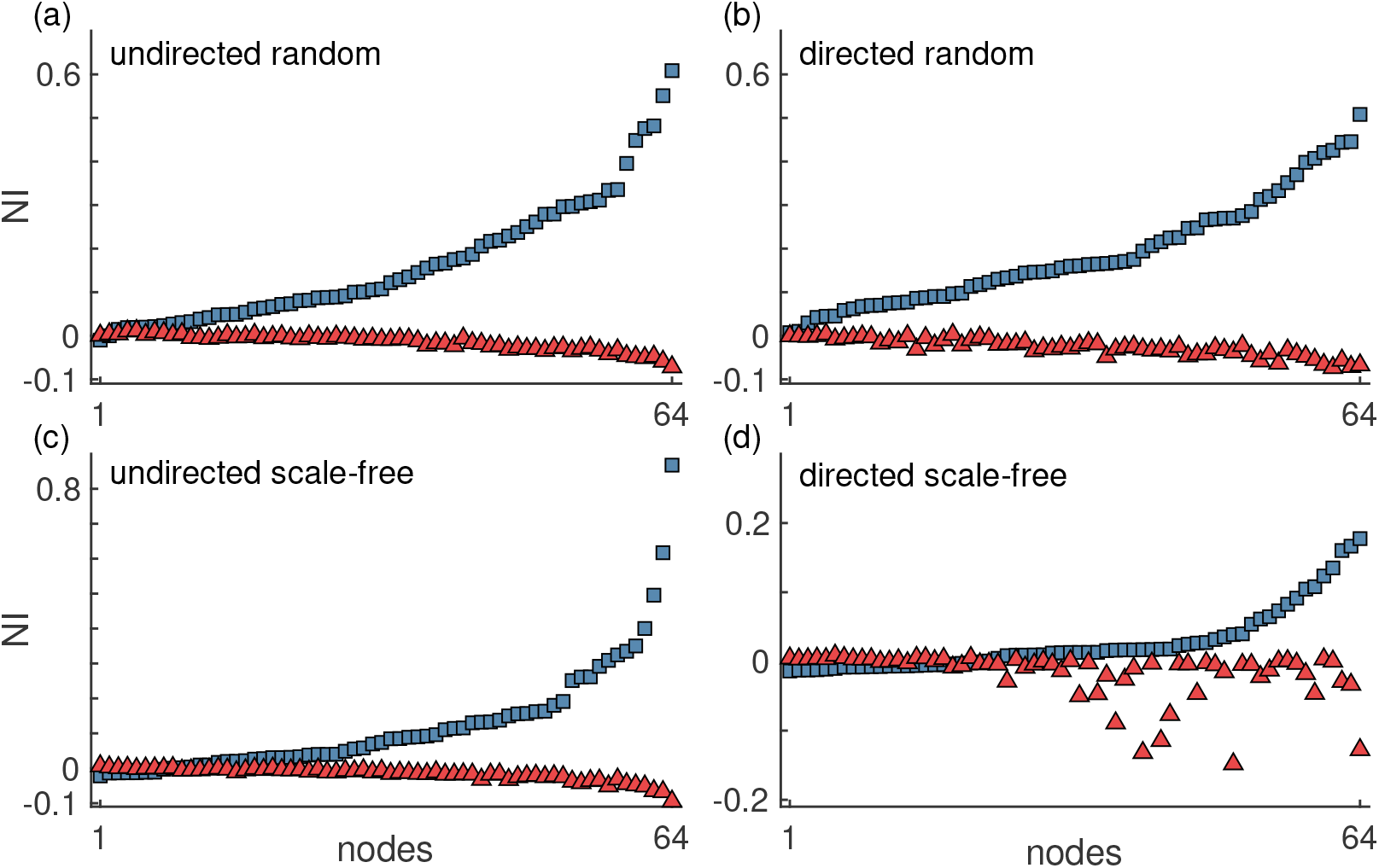
Representative NI distributions of (a) undirected random, (b) directed random, (c) undirected scale-free, and (d) directed scale-free networks using the additive and diffusive couplings. The blue squares represent the NI computed using the additive coupling and the red triangles correspond to the diffusive coupling. The nodes were sorted such that the NI grows monotonically for the additive coupling. The error bars are smaller than the symbols. The parameters *γ* and *β* were chosen such that BNI_pre_ = 0.5. All networks had mean degree *c* = 4.

**S9 Fig.**
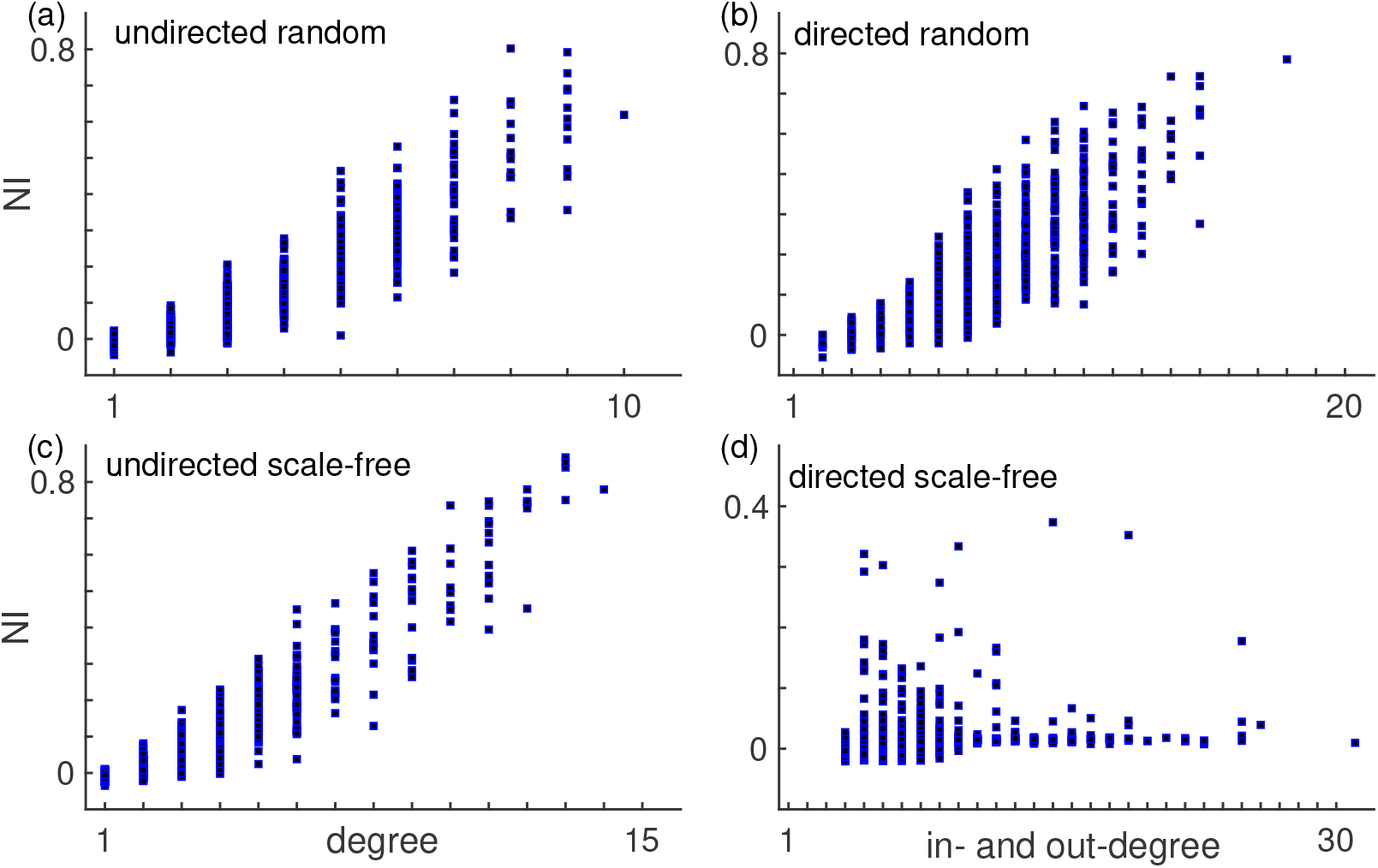
NI versus degree across different network topologies using the additive coupling and the theta model. Each panel corresponds to a different network topology: (a) undirected random, (b) directed random, (c) undirected scale-free, and (d) directed scale-free networks. In the case of the directed networks, the horizontal axis shows the sum of in- and out-degree. This figure combines the NI distributions across 10 different network realisations per network topology. The average correlation between NI and degree is 0.73 and the average p-value is 0.055. We used networks with *c* = 4.

**S10 Fig.**
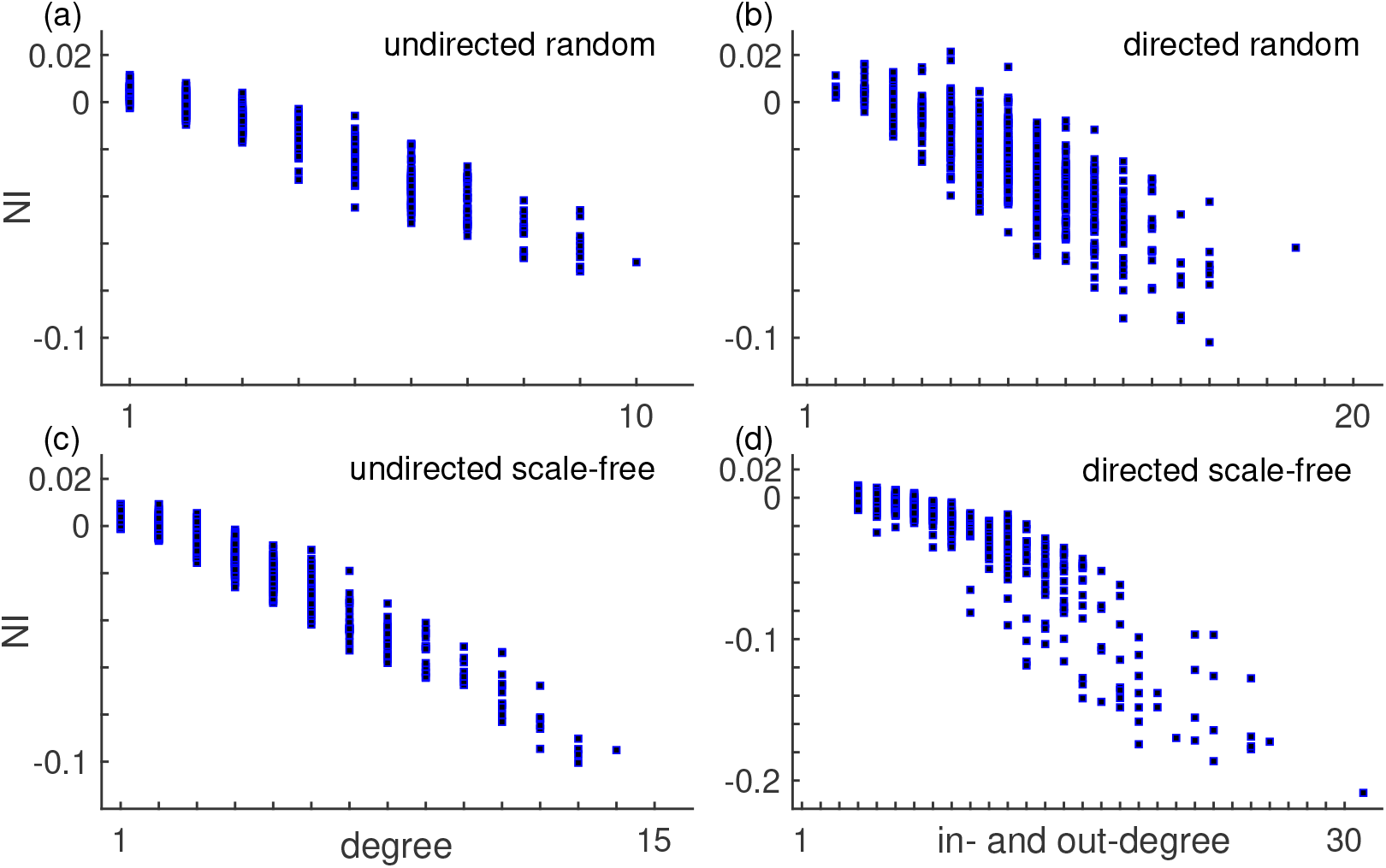
NI versus degree across different network topologies using the diffusive coupling and the theta model. Each panel corresponds to a different network topology: (a) undirected random, (b) directed random, (c) undirected scale-free, and (d) directed scale-free networks. In the case of the directed networks, the horizontal axis shows the sum of in- and out-degree. This figure combines the NI distributions across 10 different network realisations per network topology. The average correlation between NI and degree is –0.91 and the average p-value is 5.7 × 10^−16^. We used networks with *c* = 4.

**S11 Fig.**
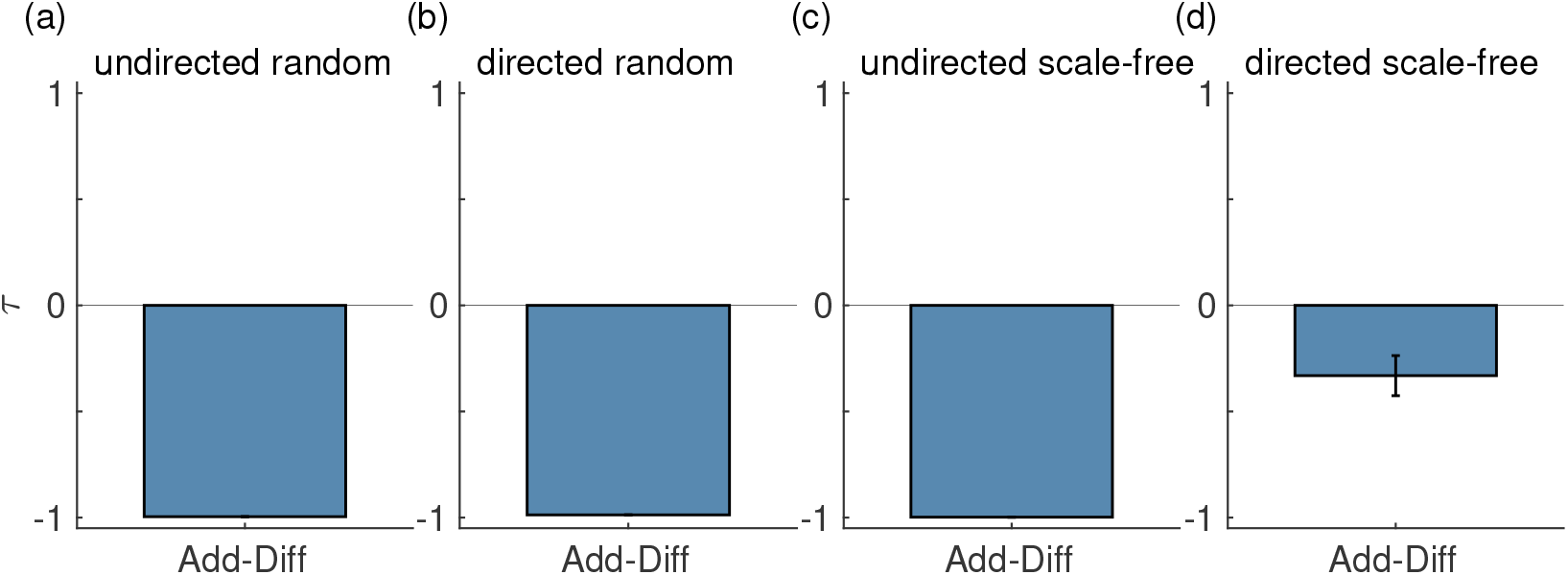
Comparison of NI distributions using the additive and diffusive couplings in the theta model. The weighted Kendall correlation rank *τ* quantifies the consistency of ordering nodes according to their NI values when using the different couplings. Different panels show the comparison between additive and diffusive coupling for different network topologies: (a) undirected random, (b) directed random, (c) undirected scale-free random, and (d) directed scale-free networks. The error bars represent the standard error across the 10 network realisations per network topology. We considered networks with *c* = 4.

Finally, S12 Fig compares the NI distributions between the theta and bi-stable models when using the same coupling function. We observe a high level of agreement in all networks, particularly when using the additive coupling. The agreement is lower in the diffusive coupling presumably because in this case the variability in NI is lower than in the additive coupling and therefore there is a higher chance of some node orderings being random.

**S12 Fig.**
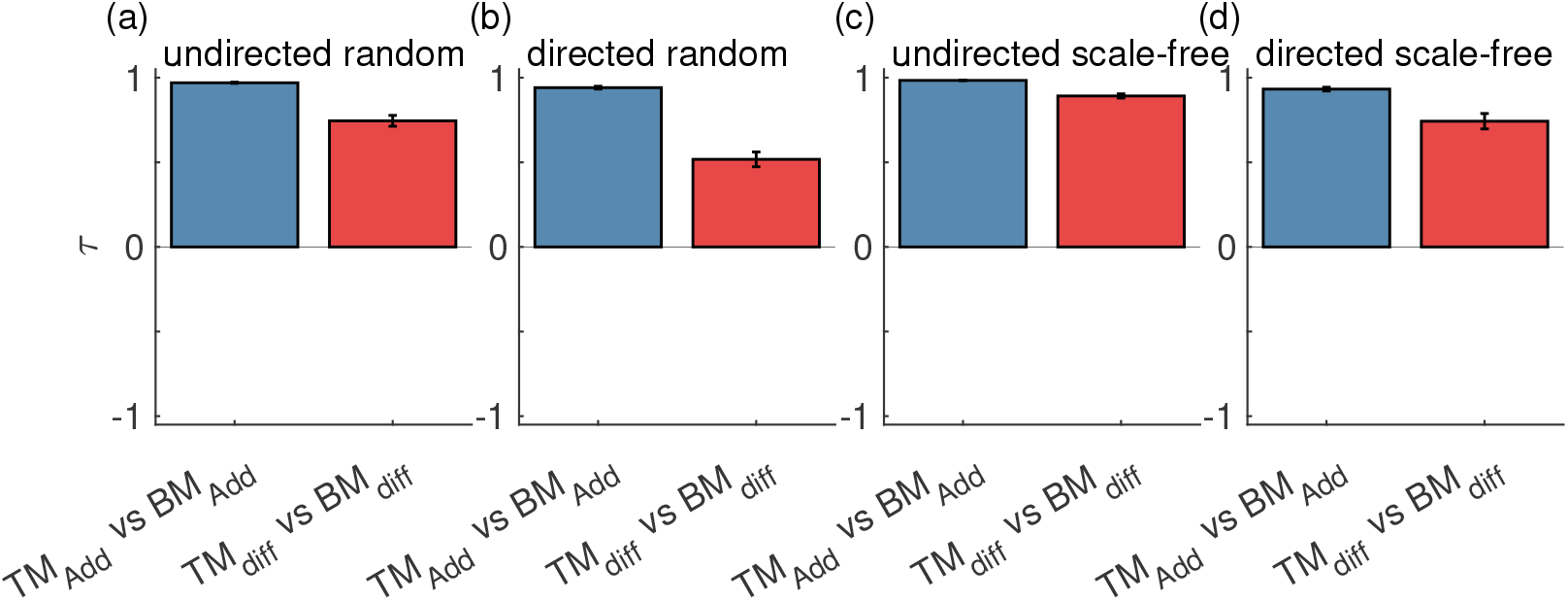
Comparison of NI distributions between the theta and bi-stable models using the additive and diffusive couplings. The blue (red) bars correspond to the comparison between the two models using both the additive (diffusive) coupling. Different panels show the comparison for different network topologies: (a) undirected random, (b) directed random, (c) undirected scale-free random, and (d) directed scale-free networks. The error bars represent the standard error across the 10 network realisations per network topology. We considered networks with *c* = 4.

## Acknowledgments

ML and JC thank John Terry and Marc Goodfellow for insightful discussions about the modelling. ML gratefully acknowledges funding from Cardiff University’s Wellcome Trust Institutional Strategic Support Fund (ISSF) [204824/Z/16/Z]. KH acknowledge the support of the UK MEG MRC Partnership Grant [MRC/EPSRC, MR/K005464/1] and a Wellcome Trust Strategic Award [104943/Z/14/Z]. KH further acknowledges a BRAIN Unit Infrastructure Award [UA05], which is funded by the Welsh Government through Health and Care Research Wales. JZ acknowledges the financial support of the European Research Council [716321]. JC gratefully acknowledges funding from MRC Skills Development Fellowship [MR/S019499/1].

## References

1. Breakspear M. Dynamic models of large-scale brain activity. Nature Neuroscience. 2017;20(3):340–352.

2. Stam CJ. Modern network science of neurological disorders. Nature Reviews Neuroscience. 2014;15(10):683–695.

3. Wilson HR, Cowan JD. Excitatory and inhibitory interactions in localized populations of model neurons. Biophysical Journal. 1972;12(1):1–24.

4. Jirsa V, Sporns O, Breakspear M, Deco G, McIntosh AR. Towards the virtual brain: network modeling of the intact and the damaged brain. Archives italiennes de biologie. 2010;148(3):189–205.

5. Hansen EC, Battaglia D, Spiegler A, Deco G, Jirsa VK. Functional connectivity dynamics: modeling the switching behavior of the resting state. NeuroImage. 2015;105:525–535.

6. Goodfellow M, Rummel C, Abela E, Richardson M, Schindler K, Terry J. Estimation of brain network ictogenicity predicts outcome from epilepsy surgery. Scientific Reports. 2016;6(1):1–13.

7. Demirtaş M, Falcon C, Tucholka A, Gispert JD, Molinuevo JL, Deco G. A whole-brain computational modeling approach to explain the alterations in resting-state functional connectivity during progression of Alzheimer’s disease. NeuroImage: Clinical. 2017;16:343–354.

8. Sanz Leon P, Knock SA, Woodman MM, Domide L, Mersmann J, McIntosh AR, et al. The Virtual Brain: a simulator of primate brain network dynamics. Frontiers in Neuroinformatics. 2013;7:10.

9. Wendling F, Bartolomei F, Bellanger J, Chauvel P. Epileptic fast activity can be explained by a model of impaired GABAergic dendritic inhibition. European Journal of Neuroscience. 2002;15(9):1499–1508.

10. Jirsa VK, Proix T, Perdikis D, Woodman MM, Wang H, Gonzalez-Martinez J, et al. The virtual epileptic patient: individualized whole-brain models of epilepsy spread. NeuroImage. 2017;145:377–388.

11. Benjamin O, Fitzgerald TH, Ashwin P, Tsaneva-Atanasova K, Chowdhury F, Richardson MP, et al. A phenomenological model of seizure initiation suggests network structure may explain seizure frequency in idiopathic generalised epilepsy. The Journal of Mathematical Neuroscience. 2012;2(1):1–30.

12. Petkov G, Goodfellow M, Richardson MP, Terry JR. A critical role for network structure in seizure onset: a computational modeling approach. Frontiers in Neurology. 2014;5:261.

13. Sinha N, Dauwels J, Kaiser M, Cash SS, Brandon Westover M, Wang Y, et al. Predicting neurosurgical outcomes in focal epilepsy patients using computational modelling. Brain. 2017;140(2):319–332.

14. Lopes MA, Richardson MP, Abela E, Rummel C, Schindler K, Goodfellow M, et al. An optimal strategy for epilepsy surgery: Disruption of the rich-club? PLoS Computational Biology. 2017;13(8):e1005637.

15. Sip V, Hashemi M, Vattikonda AN, Woodman MM, Wang H, Scholly J, et al. Data-driven method to infer the seizure propagation patterns in an epileptic brain from intracranial electroencephalography. PLoS computational biology. 2021;17(2):e1008689.

16. Hebbink J, Meijer H, Huiskamp G, van Gils S, Leijten F. Phenomenological network models: Lessons for epilepsy surgery. Epilepsia. 2017;58(10):e147–e151.

17. Junges L, Woldman W, Benjamin OJ, Terry JR. Epilepsy surgery: Evaluating robustness using dynamic network models. Chaos: An Interdisciplinary Journal of Nonlinear Science. 2020;30(11):113106.

18. Creaser J, Lin C, Ridler T, Brown JT, D’Souza W, Seneviratne U, et al. Domino-like transient dynamics at seizure onset in epilepsy. PLoS Computational Biology. 2020;16(9):e1008206.

19. Creaser J, Tsaneva-Atanasova K, Ashwin P. Sequential noise-induced escapes for oscillatory network dynamics. SIAM Journal on Applied Dynamical Systems. 2018;17(1):500–525.

20. Junges L, Lopes MA, Terry JR, Goodfellow M. The role that choice of model plays in predictions for epilepsy surgery. Scientific Reports. 2019;9(1):1–12.

21. Terry JR, Benjamin O, Richardson MP. Seizure generation: the role of nodes and networks. Epilepsia. 2012;53(9):e166–e169.

22. Doedel E, Paffenroth R, Champneys A, Fairgrieve T, Kuznetsov YA, Oldeman B, et al. AUTO-07P: Continuation and bifurcation software for ordinary differential equations. Available for download from http://indycsconcordiaca/auto. 2007;.

23. Lopes MA, Goodfellow M, Terry JR. A model-based assessment of the seizure onset zone predictive power to inform the epileptogenic zone. Frontiers in Computational Neuroscience. 2019;13:25.

24. Carterette B. On rank correlation and the distance between rankings. In: Proceedings of the 32nd international ACM SIGIR conference on Research and development in information retrieval; 2009. p. 436–443.

25. Kumar R, Vassilvitskii S. Generalized distances between rankings. In: Proceedings of the 19th international conference on World wide web; 2010. p. 571–580.

26. Newman ME. The structure and function of complex networks. SIAM review. 2003;45(2):167–256.

27. Mišić B, Sporns O, McIntosh AR. Communication efficiency and congestion of signal traffic in large-scale brain networks. PLoS Computational Biology. 2014;10(1):e1003427.

28. Rubinov M, Sporns O. Complex network measures of brain connectivity: uses and interpretations. NeuroImage. 2010;52(3):1059–1069.

29. Goh KI, Kahng B, Kim D. Universal Behavior of Load Distribution in Scale-Free Networks. Phys Rev Lett. 2001;87:278701. doi:10.1103/PhysRevLett.87.278701.

30. Albert R, Barabási AL. Statistical mechanics of complex networks. Rev Mod Phys. 2002;74:47–97. doi:10.1103/RevModPhys.74.47.

31. Lopes MA, Krzemiński D, Hamandi K, Singh KD, Masuda N, Terry JR, et al. A computational biomarker of juvenile myoclonic epilepsy from resting-state MEG. Clinical Neurophysiology. 2021;132(4):922–927.

32. Oostenveld R, Fries P, Maris E, Schoffelen JM. FieldTrip: open source software for advanced analysis of MEG, EEG, and invasive electrophysiological data. Computational intelligence and neuroscience. 2011;2011.

33. Hipp JF, Hawellek DJ, Corbetta M, Siegel M, Engel AK. Large-scale cortical correlation structure of spontaneous oscillatory activity. Nature Neuroscience. 2012;15(6):884–890.

34. Laiou P, Avramidis E, Lopes MA, Abela E, Müller M, Akman OE, et al. Quantification and selection of ictogenic zones in epilepsy surgery. Frontiers in Neurology. 2019;10:1045.

35. Cabral J, Hugues E, Sporns O, Deco G. Role of local network oscillations in resting-state functional connectivity. Neuroimage. 2011;57(1):130–139.

36. Falcon MI, Jirsa V, Solodkin A. A new neuroinformatics approach to personalized medicine in neurology: The Virtual Brain. Current opinion in neurology. 2016;29(4):429.

